# Transcription factor dynamics, oscillation, and functions in human enteroendocrine cell differentiation

**DOI:** 10.1101/2024.01.09.574746

**Authors:** Pratik N. P. Singh, Wei Gu, Shariq Madha, Allen W. Lynch, Paloma Cejas, Ruiyang He, Swarnabh Bhattacharya, Miguel M. Gomez, Matthew G. Oser, Myles Brown, Henry W. Long, Clifford A. Meyer, Qiao Zhou, Ramesh A. Shivdasani

## Abstract

Enteroendocrine cells (EECs), which secrete serotonin (enterochromaffin cells, EC) or a dominant peptide hormone, serve vital physiologic functions. As with any adult human lineage, the basis for terminal cell diversity remains obscure. We replicated human EEC differentiation *in vitro*, mapped transcriptional and chromatin dynamics that culminate in discrete cell types, and studied abundant EEC precursors expressing selected transcription factors (TFs) and gene programs. Before expressing the pre-terminal factor NEUROD1, non-replicating precursors oscillated between epigenetically similar but transcriptionally distinct *ASCL1^+^* and *HES6^hi^* cell states. Loss of either factor substantially accelerated EEC differentiation and disrupted EEC individuality; ASCL1 or NEUROD1 deficiency had opposing consequences on EC and hormone-producing cell features. Expressed late in EEC differentiation, the latter TFs mainly bind *cis*-elements that are accessible in undifferentiated stem cells and tailor the subsequent expression of TF combinations that specify EEC types. Thus, TF oscillations retard EEC maturation to enable accurate EEC diversification.

## Introduction

Specific gene programs, controlled largely by distant enhancers,^1^ underlie cell differentiation, which reflects decommissioning of *cis*-regulatory elements (CREs) that serve immature cells and recruitment of CREs that serve terminal cell states.^2^ Driving this process, tissue-restricted transcription factors (TFs) promote or sustain chromatin accessibility to ensure transcription of correct loci. Chromatin dynamics have been studied principally in differentiating embryonic or pluripotent stem cells,^3,4^ which often produce cells with fetal properties.^5–7^ Some adult epithelia, such as the skin and intestinal lining, turn over throughout life, enacting cell trajectories different from those in embryos; the determinants of rare cells within these tissues are challenging to investigate. We studied a model system for a scarce but crucial human lineage.

Enteroendocrine cells (EECs) are renewed continually in all digestive epithelia and regulate physiologic and metabolic functions from appetite, gut motility, and glycemia to microbe sensing and mucosal immunity.^8,9^ The paucity of EECs, which constitute <1% of intestinal epithelial mass, belies their physiologic importance and limits investigation of their ontogeny. Congenital EEC deficits are life-threatening in humans^10^ and mice;^11^ conversely, EEC properties can be harnessed by reprogramming, for example to secrete insulin and control hyperglycemia.^12,13^ EECs divide broadly into enterochromaffin (EC) cells, which secrete 5-hydroxytryptamine (5HT, serotonin) and various non-EC cells, each of which produces a dominant or at most a few of the >20 known enteric peptide hormones,^14^ such as glucagon-like peptide 1 (GLP-1 – an incretin), ghrelin (GHRL – an orexigen) or somatostatin (SST – an antagonist of many hormones); GLP-1 agonists are used to treat type 2 diabetes and obesity.^8^ Because EECs sense and respond to local signals and many peptide hormones oppose one another, limiting the array of hormones in any EEC is a physiologic adaptation. Some EECs modify their hormonal output as they migrate along intestinal villi,^15^ in response to local cues,^16^ but the basis *a priori* for hormone-expressing potential is not well appreciated. Knowing that basis has both fundamental and medical value.

EECs originate in secretory daughters of intestinal stem cells (ISCs). NEUROG3, a TF that appears transiently in early EEC progenitors, is necessary and sufficient to drive mouse and human EEC differentiation,^10,11,17–19^ but how downstream TFs^20–24^ interact with chromatin and with each other to generate discrete mature EEC types is unknown. Bulk^25^ and single-cell RNA sequencing (scRNA-seq) of mouse^26–28^ and human^29^ EECs has highlighted species and cell similarities and differences, but CRE dynamics remain unknown. One limitation is that even after perturbing culture conditions^30^ or forcing NEUROG3 expression in organoids cultured from induced pluripotent stem cells^19^ or tissue biopsies,^29^ it is difficult to capture the full spectrum of EEC differentiation.

By activating *NEUROG3* transiently in 2D human ISC cultures, we triggered full and efficient EEC differentiation, and we examined stepwise modulation of enhancers and genes at single-cell (sc) and single-nucleus (sn) resolution. We find that stages between the earliest EEC precursors and *NEUROD1^+^* pre-terminal EECs constitute an interval of limited TF and chromatin dynamics. Within that watershed interval, cells oscillate between epigenetically similar but transcriptionally distinct cell states marked by expression of basic helix-loop-helix (bHLH) TFs ASCL1 and HES6. Depletion of these TFs revealed that the oscillatory phase substantially *restrains* EEC differentiation and enables accurate subsequent diversity. Although ASCL1 and NEUROD1 first appear days after NEUROG3 activation, most of their binding occurs at genomic DNA where chromatin is accessible even in ISCs and their substantial effects on EEC fidelity and maturation occur with little chromatin perturbation, reflecting dysregulated expression of downstream TFs. Thus, ASCL1, HES6, and NEUROD1 accurately sculpt the TF repertoire and combinations that distinguish EC from non-EC cells and various non-EC cell types from one another. This study of an adult human lineage reveals the ordered cellular trajectory for proper diversification of medically important cells, providing molecular insights that could be useful to engineer EECs for therapy.

## Results

### Human EEC differentiation and divergence in 2D culture

To culture human ISCs (hISCs), we generated 3D organoids from small intestinal crypts, then propagated cells over mitotically inactive embryonic mouse fibroblasts.^31^ In 2D cultures, hISCs proliferated in colonies, expressed *SOX9*, *LGR5*, *OLFM4* and other ISC markers, preserving regional identity over dozens of passages (Figures S1A and S1B). Because transient NEUROG3 expression specifies mouse and human EECs *in vivo* and *in vitro*^10,18,19,32^ we engineered hISC lines to express tamoxifen (Tam)-inducible mouse NEUROG3 and mCherry (mCh, Figure 1A). Independent ileal hISC^NEUROG^^3^ lines treated with Tam ceased proliferation within 2 days and colonies gradually dispersed; by 6 days, >95% of mCh^+^ cells expressed Chromogranin A (CHGA, an EEC marker) and subsets of cells expressed 5HT (serotonin), SST or GLP1 (Figures S1C and S1D). Two months after we delivered Tam-treated cells into the portal veins of immunodeficient mice, recipient livers carried numerous mCh^+^ 5HT^+^ GLP1^+^ KI67-cell clusters (Figure S1E). Thus, hISC^NEUROG^^3^ lines yield stable and essentially pure EECs.

**Figure 1.**
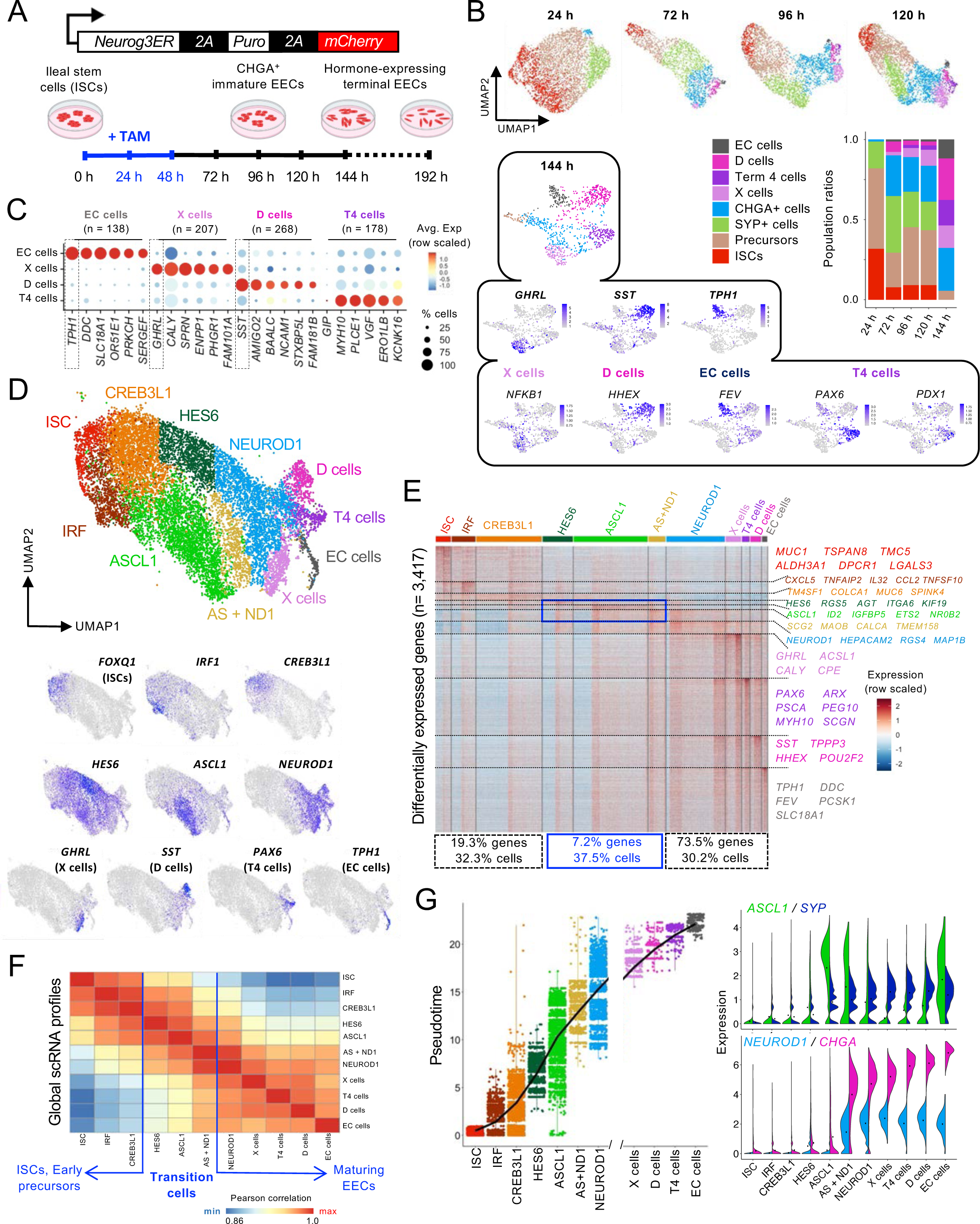
Advanced differentiation of human EEC types *in vitro*. (A) Overview of study design. A polycistronic EF1α-*Neurog3^ER^-Puro^R^-mCherry* construct was introduced stably into 2D human ISCs for constitutive expression of NEUROG3^ER^ (inducible with 4-OH tamoxifen, Tam) and mCherry. Exposure of hISC^NEUROG^^3^ cells to Tam for 48 h initiates EEC differentiation, which occurs over ∼6 days. (B) scRNA-seq derived UMAP clustering of mCherry^+^ cells isolated by FACS, 24 h (n=3,240 cells), 72 h (n=1,497), 96 h (n=2,727), 120 h (n=3,449), and 144 h (n=1,402) after the start of Tam treatment. Cell clusters were identified using markers *FOXQ1* (ISCs), *SYP* and *CHGA* (generic for EECs), *GHRL* (X cells), *SST* (D cells), *TPH1* (EC cells), and *PAX6* (T4 cells). Early precursors express reduced levels of ISC markers and low levels of *SYP*. Cell proportions across time show progressive decline of ISCs, with gradual increase in hormone-expressing cells. Feature plots at 144 h for hormone-encoding and restricted transcription factor (TF) genes are shown in the box: *GHRL* and *NFKB1* for X cells, *SST* and *HHEX* for D cells, *TPH1* and *FEV* for EC cells, and *PAX6* and *PDX1* for T4 cells. (C) Relative expression of unique marker genes enriched in each EEC type, where *TPH1*, *SST* and *GHRL* delineate EC, D, and X cells, respectively. (D) Integrated merged UMAP of scRNA-seq derived cells (n=12,315 cells) from 24 h, 72 h, 96 h, 120 h and 144 h after NEUROG3 induction. Cell clusters are defined by their nearest neighbors and selected TFs enriched in each state are shown in the feature plots below. (E) Genes (n=3,417) differentially expressed across 11 cell clusters identified from scRNA-seq and the proportions of cells and differentially expressed genes in early precursors, transition cells (blue boxes), and *NEUROD1^+^*maturing EECs. The top differential genes in each cluster are listed to the right. (F) Pearson correlation of global transcriptional profiles for the scRNA-seq clusters. ISCs and early precursors on the one hand, and maturing EECs on the other, are more different compared to transition cells. (G) Monocle3-derived trajectory of cell clusters identified by scRNA-seq, showing progression of cells along a pseudotime axis from hISC^NEUROG^^3^ to terminal EEC types. Right: *ASCL1*, which coincides with the onset of *SYP* expression, precedes *NEUROD1*, which coincides with the onset of *CHGA* expression, in pseudotime. See also Figures S1 and S2.

Two independent hISC^NEUROG3^ lines showed progressive mRNA alterations associated with NEUROG3-induced EEC differentiation. To exclude Tam effects unrelated to NEUROG3, we generated additional cells expressing doxycycline (Dox)-inducible NEUROG3. After Tam or Dox exposure, we observed quantitative changes (>4-fold between any two time points, *q* <0.05) in ∼4,000 genes (Figures S1F to S1H and Table S1). Transcripts enriched in untreated cells serve stem-cell functions and their attenuation after NEUROG3 activation gave way first to an mRNA response associated with interleukin, interferon γ and STAT signaling (Figures S1G to S1I), followed by expression within 96 h of markers *SYP* and *CHGA*, and later of multiple hormones (Figures S1G to S1J). Immunoblots confirmed STAT signaling (Figure S2A). Presence of *SST*, tryptophan hydroxylase (*TPH1*), ghrelin (*GHRL*), glucose-dependent insulinotropic polypeptide (*GIP*) and other transcripts (Figures S1J and S2B) signified EEC differentiation encompassing at least delta (D), EC, X, and K cells, respectively. More than 300 TF transcripts changed >4-fold between any 2 days (*q* <0.05, Figure S2C), including known EEC-associated TFs *NKX2-2*, *NEUROD1*, *RFX6*, *PAX4*, *PAX6*, and *ARX* (Table S2). These included 75 of the 172 TF genes (43.60%) identified in mouse^28^ and 15 of 22 identified in human^29,33^ EECs. Thus, hISC^NEUROG3^ cells allow temporal monitoring of EEC differentiation; major EEC types were sufficiently specified within 144 h of TAM exposure to investigate lineage diversification (Figure S2D).

We therefore captured scRNA profiles of hISC^NEUROG3^ cells collected 24 h (n=3,240), 72 h (n=1,497), 96 h (n=2,727), 120 h (n=3,449), and 144 h (n=1,402) after Tam exposure. Graph-based clustering analysis and display of cell populations by uniform manifold approximation and projection (UMAP)^34^ (Figures 1B and S2E) indicated asynchronous EEC differentiation, with *SYP^+^ and CHGA^+^* cells and their precursors arising continually and a few ISCs persisting up to 120 h (Figures 1B and S2E). Of the 4 cell clusters present at 144 h and representing mature EEC sub-types, three expressed specific markers and a predominant hormone (Figure 1B-C), signifying EC (*TPH1*/serotonin dominant), X (*GHRL* dominant), and D (*SST* dominant) cells. The fourth cluster, “terminal 4” (T4), expressed low levels of various hormones (e.g., GIP – Figure S2F) and abundant *PAX6* and *PDX1* (Figure 1B), TFs implicated in mouse K and L cells.^28,35^ T4 thus represents either cells fated in aggregate for K, L, and other peptide-secreting identities or a composite EEC producing multiple hormones,^26,36,37^ separate from X or D cells. Terminal *TPH1^+^* EC cells were distinct from peptide-producing (non-EC) cells and mRNA signatures defined each EEC type (Figures 1C, S2G, S2H and Table S3).

Integration of scRNA data from all time points delineated 11 cell clusters, which we name as ISCs, mature EEC types, or by the TF gene most enriched in expression (Figure 1D and Table S4). Differential gene expression correlated with the time after TAM exposure (Figure S2E) and with the mRNA trajectories and dominant biological processes revealed in bulk mRNA profiles (Figures 1E and S3A), including IFN and interleukin signaling in early EEC precursors (Figure S3B). Among 3,417 differentially expressed genes (ln fold change >0.2, *P_adj_* <0.05, detected in ≥25% of cells within an enriched cluster – Table S3), 19.3% were enriched in ISCs or early precursors and 73.5% were enriched in late cells, starting in a cluster dominated by *NEUROD1* (Figure 1E). The remaining 7.2% of differential genes were enriched in abundant transitory populations (37.5 % of all cells) characterized by *ASCL1* (2,822 cells) or high *HES6* (1,149 cells) expression. Global scRNA profiles of these two clusters placed them in a transcriptional watershed between early EEC precursors and *NEUROD1^+^*maturing EECs (Figure 1F).

### Transcriptional, TF, and chromatin trajectories in human EEC differentiation

Pseudo-temporal (PT) analysis of EEC differentiation using Monocle3 (Ref. ^38^) positioned the cell clusters between ISCs and mature EECs in the sequence IRF è CREB3L1 è HES6 è ASCL1 è NEUROD1, followed by emergence of cells recognized as X, D, T4, and EC from a parental *NEUROD1*-dominant state (Figures S3C and 1G). Immunoblots of bulk cultures verified (Figure S3D) that *ASCL1* preceded *NEUROD1* expression and then declined, with a transient intermediate state expressing both TFs (Figure 1G). After transient expression in EEC precursors, *ASCL1* was attenuated in all terminal cells other than the *TPH1^+^* EC fraction, whereas *NEUROD1* persisted in all mature cells (Figure 1G).

Among the 287 TF genes differentially expressed between any two clusters along the PT trajectory (ln fold-difference >0.2, *P_adj_* <0.05), most appeared after *ASCL1* levels had fallen (Figure 2A) and 38 were detected at levels sufficient for meaningful interrogation after genetic perturbations (min. average expression >1.0 count per cell) and selectively enriched (fraction of cells expressing in *that* cluster >0.45, and difference from fraction of cells expressing in *other* clusters >0.2). Across mature EECs, these 38 TF genes appeared in various combinations, with few TFs exclusive to one cell type. For example, *ISL1*, *MAFB, PBX3* and *ARX* appear in various non-EC cells and *FEV*, *DACH1, RFX3, POU2F2, PBX1*, *PAX6,* and *ASH1L* appear in EC and some non-EC cells (Figures 2B and S3E). Thus, distinct waves of TF expression precede and follow a state defined by *ASCL1* dominance, onset of *SYP* and *SOX2* expression, absence of hormone mRNAs, and limited TF dynamics. A disproportionately large fraction of cells occupied this watershed state, which was distributed more widely along the PT continuum than any other (Figure 2A), suggesting that precursors might spend more time in this state than in others.

**Figure 2.**
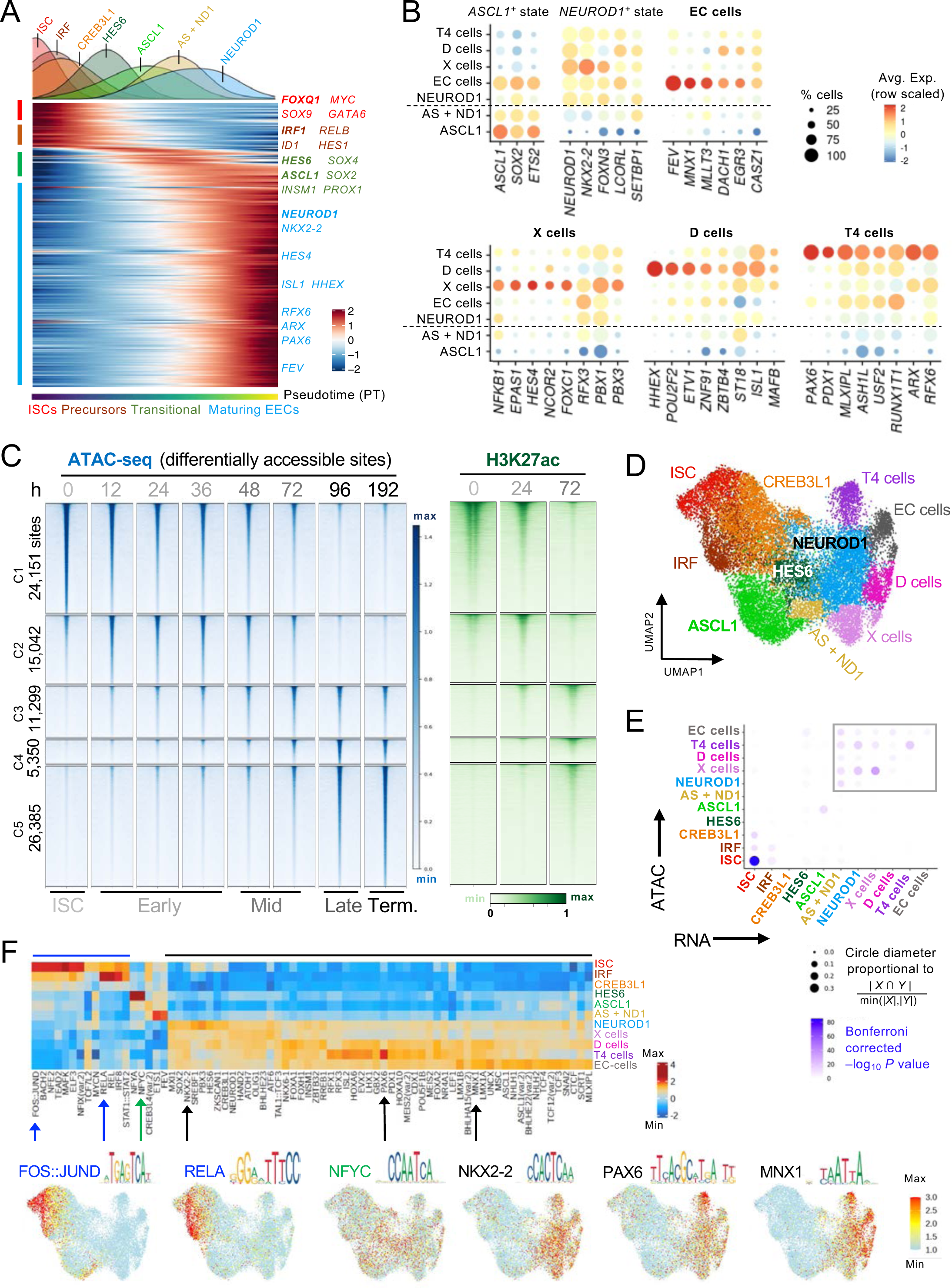
Dynamics of TF genes and chromatin access during EEC differentiation. (A) Differential expression of 287 TF genes in single cells along the pseudotime trajectory, showing a gradual decline in levels of 53 TFs in ISCs (red) and early precursors (brown), transient expression of few TFs in transitional cells (green), and late expression of 214 TFs in maturing EECs (blue) after *ASCL1* levels fall. (B) Relative expression of TF genes expressed differentially and at sufficient levels in the *ASCL1^+^* state and beyond. Rare TFs are restricted to one EEC type; most are expressed in different amounts in overlapping populations. (C) Differentially accessible genomic sites identified from bulk-ATAC seq analysis of mCherry^+^ hISC^NEUROG^^3^ cells isolated at the specified times after Tam exposure (NEUROG3 induction). Right, H3K27ac Chip-seq signals at the specified times after Tam exposure at sites showing differential chromatin access. (D) Integrated merged UMAP of snATAC-seq derived cells (n=18,954) from 24 h, 72 h, 96 h, 120 h and 144 h after Tam induction. The 11 clusters correspond to scRNA-seq defined cell clusters (Figure 1D) by transfer of gene anchors. (E) Overlap among clusters from stand-alone scRNA and snATAC modalities, quantified by the intersects between genes enriched for expression in each cluster (rows: scRNA data) and the nearest site of accessible chromatin in the corresponding cells (columns: snATAC data). Circle sizes represent the degree of mRNA:ATAC peak correspondence for expressed genes and the color scale represents Bonferroni-corrected *P* values from a hypergeometric test. High concordance is evident in untreated hISC^NEUROG^^3^ (bottom left) and in *NEUROD1^+^* maturing EECs (boxed area, top right). (F) TF consensus motifs enriched at chromatin sites selectively accessible in each pseudobulk snATAC cell cluster, displayed as deviations (row scaled) from expected motif abundance. Accessibility signals (z scores from chromVAR) for representative enriched motifs (arrows) are projected below onto the integrated UMAP: FOS/JUND and RELA for ISCs and early precursors, NFYC for the transition state, and NKX2-2 (which appears in all EECs after *NEUROD1*), PAX6 (enriched in T4 cells) and MNX1 (enriched in terminal EECs including EC cells) for maturing EECs. See also Figures S3 and S4.

Because gene activity requires accessible chromatin at CREs, especially enhancers,^39^ we used the assay for transposase-accessible chromatin (ATAC)^40^ to examine dynamic chromatin access during EEC differentiation; replicate samples on different days after Tam exposure gave concordant data (Figure S3F). More than 80,000 enhancers (regions >2 kb from transcription start sites) were modulated (>4-fold change, *q* <0.05) between any 2 days, broadly representing early (12 to 36 h post-Tam), mid (48 to 72 h), and late (96 h) transitions from ISCs to mature (192 h post-Tam) EECs (Figure 2C). H3K27ac, an active histone mark we assessed by chromatin immunoprecipitation (ChIP-seq) in untreated cells and 24 h or 72 h after Tam exposure, coincided with this modulation of chromatin access (Figure 2C). Loci active in ISCs (e.g., *FOXQ1*, *MYC*) reduced chromatin access, while others (e.g., *SERPINA4*) gained access within 12 h (Figure S3G). The profile of open chromatin in terminal EECs appeared by 96 h (Figure 2C) and enhancer dynamics were correlated with temporal mRNA profiles (Figure S3H). Furthermore, DNA sequence motifs enriched at stage-selective CREs matched the onset or peak expression of the corresponding TFs, e.g., ETS, IRF and NF-κB motifs at the early transition, SOX, ASCL1 and NEUROD1 motifs at subsequent transitions, and culminating in motifs for NKX2-2, RFX, ISL1 and other TFs that dominate in terminal EECs (Figure S4A). Thus, transcriptional and chromatin dynamics together reveal successive waves of TF expression and CRE activation in human EEC differentiation.

To further resolve dynamic chromatin accessibility, we profiled open chromatin in single cells isolated 24 h (n=1,459), 72 h (n=1,543), 96 h (n=3,652), 120 h (n=5,330), and 144 h (n=6,970) after Tam treatment (Figure S4B). Sites identified by bulk and scATAC overlapped significantly (Figure S4C) and transfer of gene anchors from scRNA allowed us to define the corresponding cell clusters across an integrated UMAP of open chromatin profiles in single cells from all days (Figure 2D). Chromatin open selectively in ISCs or in maturing *NEUROD1^+^* cells correlated with genes expressed in the corresponding scRNA clusters; however, mature EECs resembled each other in ATAC and mRNA profiles more than preceding cell states (Figure 2E). We used Monocle3 to order UMAP cell clusters in PT,^34,38^ anchored by marker loci expressed in each cluster (Figure S4D). In agreement with the resulting PT order, we observed decommissioning of ISC enhancers followed by waves of new CRE accessibility (Figures S4E and S4F). DNA motifs for the TFs expressed exclusively in any scRNA cluster, e.g., FOSL2, TEAD, and RELA (early) and NKX2-2, PAX6, MNX1, and others (late), were enriched in the corresponding scATAC cluster (Figure 2F). Starting with *NEUROD1^+^* cells, however, mature EECs, shared large fractions of enhancers and DNA motifs (Figure 2F), indicating that discrete EEC properties arise within a shared program of accessible chromatin. Notably, various TF motifs were highly enriched in ISCs, EEC precursors, and maturing *NEUROD1^+^*cells, but almost none were enriched in the 3,827 interim cells (20.6% of the total) characterized by *ASCL1* or high *HES6* expression (Figure 2F). This finding reinforces the idea that the latter cell states represent a watershed between decommissioning of ISC-restricted and recruitment of EEC-specific CREs.

### Atypical cell dynamics precede generation of NEUROD1^+^ EEC precursors

Because integrating stand-alone scRNA- and scATAC-seq has theoretical limitations^41^ and to enhance confidence in the above PT inferences, we used a multi-ome platform to map RNA expression and chromatin access simultaneously in 16,898 nuclei isolated 24 h (n=7,055 cells), 72 h (n=2,742), 120 h (n=3,653), and 144 h (n=3,448) after Tam exposure (Figure S5A). To map EEC ontogeny in PT, we applied MIRA, a probabilistic multimodal model for integrated regulatory analysis based on topic representation of cell states and the regulatory potential of individual loci.^42^ This unsupervised approach, which eschews prior definition of “nearest neighbor” clusters, nevertheless allowed us to recognize cell states by their similarities to those revealed by stand-alone scRNA-seq, i.e., hormone-expressing terminal EECs and transitory cells dominated by *IRF1*, *HES6*, *ASCL1*, etc. (Figures 3A and S5B). Furthermore, integrated consideration of chromatin accessibility features, DNA motif enrichment, and RNA expression in individual cells over the MIRA-derived trajectory resolved distinct temporal regulatory modules (Figure 3B). After NEUROG3 induction, ISC-specific *cis*-elements became inaccessible, followed by transient chromatin accessibility at CREs associated with interferon signaling, followed by new and sustained access leading to activation of terminal EEC genes. From the *NEUROD1^+^* precursor state, both RNA and ATAC topics indicated that four distinct *cis*-regulatory genomic profiles, corresponding to X, D, K and T4 cells, arise from combinatorial TF binding at identifiable DNA motifs (Figure S5C). Notably, cell division was restricted to ISCs and small fractions of *IRF^+^* or *CREB3L1^+^* early EEC precursors, but absent in *HES6^hi^* or *ASCL1*- and *NEUROD1*-expressing cells (Figure S5D), consistent with known potent NEUROG3 activity in cell cycle repression.^43^ Thus, combined consideration of mRNA and chromatin dynamics in single nuclei verifies EEC divergence, long after mitosis, from a common *NEUROD1^+^* precursor (ISCèEC and ISCèT4 cell trajectory examples in Figure 3C).

**Figure 3.**
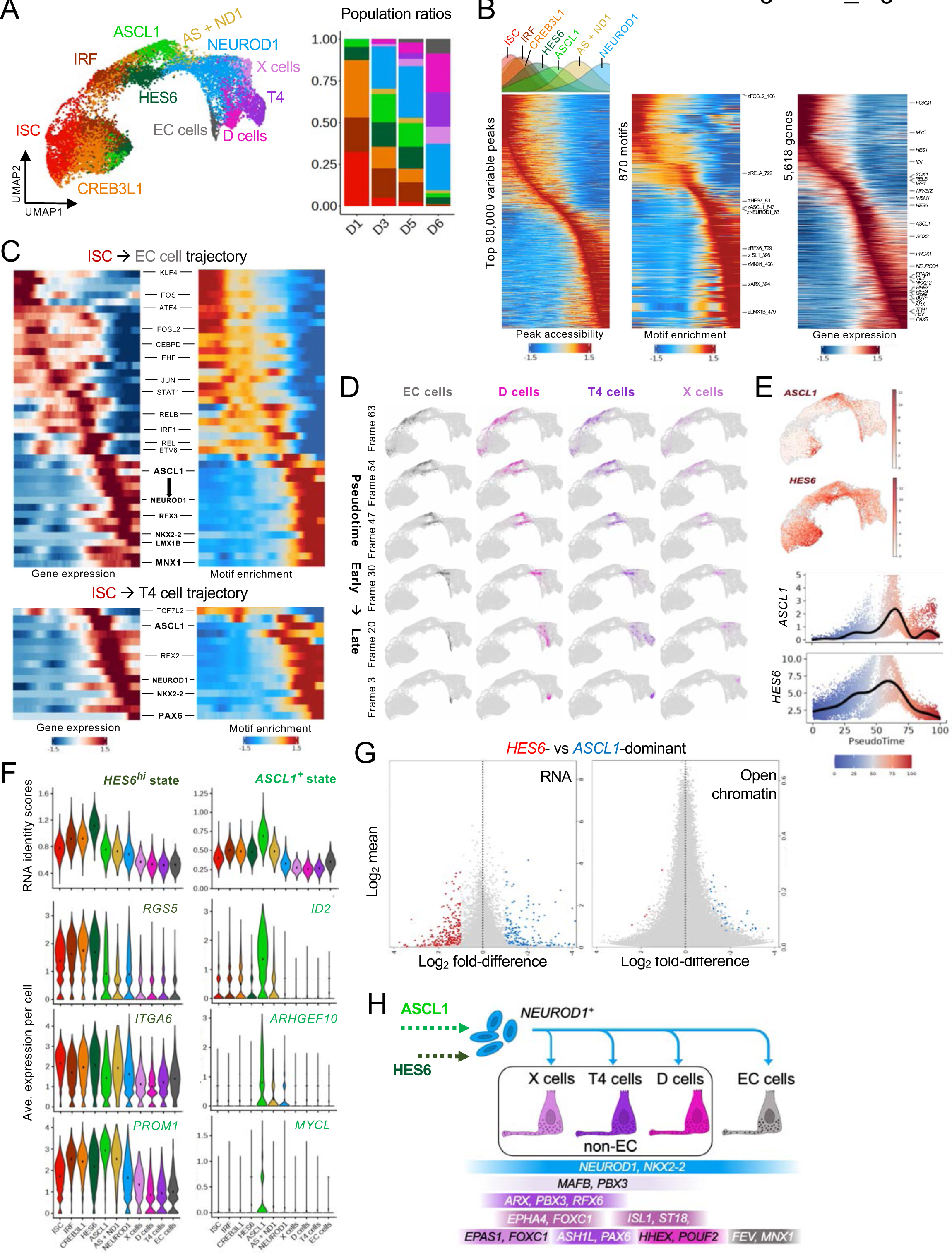
Trajectory of cellular states culminating in mature EEC derivatives. (A) Multimodal model for integrated regulatory analysis (MIRA)-derived UMAP of snMultiome-seq (RNA + ATAC) of cells merged from 24 h (n=7,055 cells), 72 h (n=2,742), 120 h (n=3,653) and 144 h (n=3,448) after NEUROG3 induction. Cell positions in pseudotime, defined by topic representation in a probabilistic model, are represented in a k-nearest neighbor (KNN) graph. Groups of cells with common properties are labeled based on their correspondence (Figure S5B) to gene anchors from previously defined scRNA-seq clusters (Figure 1E). (B) MIRA-derived pseudotime trajectories of peak accessibility (ATAC component), motif enrichment relative to the genome background, and TF gene expression (RNA component) from snMultiome data. Note the high concordance between differential TF genes and chromatin access at genomic sites that carry the corresponding sequence motifs. (C) Correlation of TF gene expression (left) with TF sequence motifs (right, deviations from expected abundance, chromVAR) across the ISC è EC and ISC èT4 cell trajectories. (D) Pseudotime trajectories in the KNN graph representing differentiation of each terminal EEC cluster, in reverse pseudotemporal flow of cells from ISCs to mature EECs. Selected frames from Supplemental Videos SV1 to SV4 reveal the shared ancestry of each EEC type in *both ASCL1*-dominant and *HES6*-dominant precursors. (E) *ASCL1* and *HES6* expression projected on the MIRA-derived KNN graph. Pseudotemporal graphs below show earlier onset of *HES6* than of *ASCL1* expression and concomitant peaking of both genes. (F) Expression of *HES6^hi^*and *ASCL1^+^* transcriptional state signatures (from Table S3, top) and individual state-enriched genes (below) in each cell cluster along the pseudotime trajectory. (G) Differential RNA expression (left) and chromatin accessibility (right) in *HES6*- and *ASCL1*-dominant EEC precursor cell states, revealing abundant RNA but little chromatin variability. Y-axes: average normalized RNA or ATAC signals, X-axes: log_2_ fold difference between the two states. Red and blue dots represent genes/genomic sites that are, and grey dots represent those that are not, significantly altered (*q* <0.001, log_2_ fold-difference |>1|). (H) Model for EEC cell states along the differentiation trajectory. Selected markers defining each terminal cellular state are represented at the bottom. See also Figure S5.

This shared *NEUROD1^+^* node revealed unusual antecedents: all cells entered it by passing through *both ASCL1*-dominant *and HES6*-dominant states (Figure 3D and Supplemental Videos SV1 to SV4). While *ASCL1* appeared *de novo* during this period in the PT trajectory, *HES6* was present at low levels *ab initio* and peaked coincident with *ASCL1* (Figures 3E and S5E). Thus, early EEC precursors seem to branch in two paths, which overlap in pseudotime and have fewer enriched markers than other states (Table S3). Notably, the 43 genes enriched in *HES6^hi^* cells together distinguish those cells marginally, while the 54 genes enriched in the *ASCL1^+^* cluster distinguish it (Figure 3F) at least as well as any mRNA signature distinguishes mature EEC types (Figure S2G). Indeed, although *HES6^hi^* mapped as a discrete cluster, its individual markers are modestly enriched over other states, in contrast to *ASCL1*-synexpressed genes, which are specific to *ASCL1^+^*precursors and largely absent even from *ASCL1*-expressing EC cells (Figures 3F and S5E). Some *ASCL1^+^* precursor-enriched genes were expressed broadly (e.g., *PROM1* – Figure 3F) or persisted in ECs (e.g., *ETS2* – Figure S5E) but overall, *ASCL1^+^* cells constituted a distinct precursor state, while *HES6^hi^* represents the end of a PT continuum extending from *IRF^+^* and *CREB3L1^+^* cells (Figure 3F).

One possibility is that *ASCL1* expression signifies early EC specification while non-EC cells take a parallel, *HES6*-dominant path. In this case, EC and non-EC cells would show different trajectories preceding the *NEUROD1^+^* node, *ASCL1^+^* cell numbers would match those of EC cells, and the *NEUROD1^+^* state would represent a mix of specified EC and non-EC cells. However, *all* mature EEC origins traced to *both ASCL1*^+^ and *HES6^hi^* states (Figure 3D and Supplemental Videos SV1 to SV4), *ASCL1^+^* cell numbers always greatly exceeded the EC output (Figures 1E and 2F), and *NEUROD1^+^* cells were uniform, lacking signs of EC or non-EC specification (Figure S5B). Of further note, although ∼400 mRNAs distinguished *ASCL1^+^* from *HES6^hi^* cells, open chromatin differed only at selected loci in *ASCL1^+^* cells (Figure 3G). Thus, *NEUROD1^+^* cells arise from transcriptionally distinct but epigenetically related precursor states before different TF gene combinations resolve terminal EEC types (Figure 3H).

### TF oscillation in the EEC differentiation watershed

*Ascl1* is expressed in mouse neural progenitors and *Ascl1*^-/-^ mice lack substantial neuronal populations: autonomic and olfactory neurons,^44^ serotonergic brain and gut neurons,^45,46^ retinal bipolar cells^47^. In mouse neuronal and oligodendrocyte progenitors, ASCL1 and HES-family TFs oscillate in alternating phases before cells differentiate.^48,49^ HES1 represses *Ascl1* transcription in mouse neuronal progenitors (NPs)^50^ and HES6 counters that repression.^51^ Optogenetically driven ASCL1 oscillation promotes NP cell replication and, conversely, sustained non-oscillatory expression enhances neuronal differentiation.^48^ These precedents raised the possibility that *ASCL1^+^*and *HES6^hi^* EEC precursors represent oscillating, and not parallel or sequential, cell states. Indeed, scRNA-seq data for genes enriched in each precursor, such as *ASCL1* or *HES6* themselves and *KLF13*, revealed high unspliced RNA (signifying active transcription) in one state and measurable levels of spliced, steady-state transcripts in the other (Figure 4A); this reciprocity suggests oscillating rather than parallel or sequential cell states. Despite the typically lower contrast between spliced and unspliced transcripts in snMultiome libraries, which lack cytoplasmic RNAs, unspliced *HES6* and *KLF13* transcript residuals were confined to HES6^Hi^ cells, while the spliced form extended into *ASCL1^+^*cells (Figure S5E).

**Figure 4.**
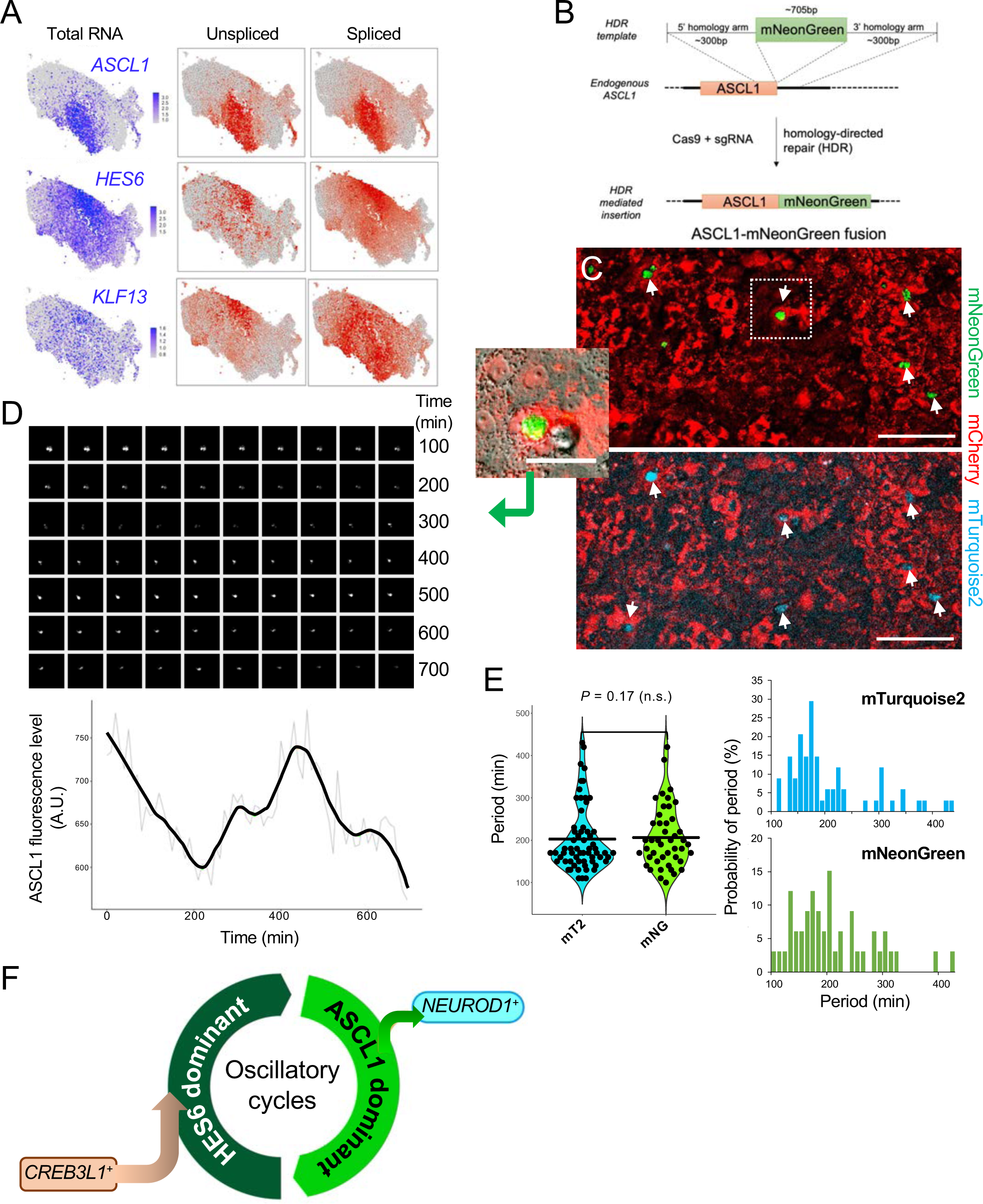
ASCL1 oscillation during human EEC differentiation. (A) Levels of total, unspliced and spliced *ASCL1*, *HES6*, and *KLF13* transcripts projected onto the integrated UMAP from stand-alone scRNA-seq. Unspliced *HES6* and *KLF13* transcripts (reflecting ongoing gene activity) are restricted to the *HES6*-dominant state, compared to the spliced forms (reflecting prior gene activity), which are present in a broad domain that encompasses *ASCL1*-dominant cells. (B) Strategy to generate fluorescence-tagged *ASCL1* by homology-directed gene editing in hISC^NEUROG^^3^ cells. (C) Representative fluorescence images of nuclear ASCL1-mNeonGreen (mNG) and ASCL1-mTurquoise2 (mT2) fusion proteins 72 h after Tam treatment to induce NEUROG3 activity (scale bar 100 µm). The mNG^+^ cell boxed in the top panel is magnified to the left (scale bar 50 µm). (D) Time-lapse images (top, 10-min intervals) and quantification of nuclear ASCL1 fluorescence (bottom, raw data and smoothed curve) from the representative mNG^+^ cell magnified in panel C. Additional examples are shown in Figure S5H. (E) Distribution of ASCL1 oscillation periods observed with mNG (n=20 nuclei) and mT2 (n=21 nuclei) reporters. Periods were similar with either label. n.s., not significant by paired t-test. (F) A phase of oscillation between *ASCL1*- and *HES6*-dominant states separates *CREB3L1^+^ HES6^+^* early EEC precursors from *NEUROD1^+^* late EEC precursors. See also Figure S5.

To test directly for ASCL1 oscillation, we used Cas9/CRISPR homologous recombination to insert 3’ mNeonGreen (mNG) or mTurquoise2 (mT2) fluorescent tags^52^ into the *ASCL1* locus in hISC^NEUROG3^ cells (Figure 4B). After exposure to mNG and mT2 cDNAs in equal proportion, up to 5% of cells showed nuclear fluorescence with one or both labels (Figure 4C) and DNA sequencing verified in-frame recombination (Figure S5F – a parallel effort for the *HES6* locus was inefficient). Starting 72 h after Tam exposure, and over the ∼12 h that cells stayed healthy under observation, we used time-lapse fluorescence imaging at 10-min intervals to monitor those expressing ASCL1 fusion proteins. Nuclear signals were oscillating, with co-expression of mNG- and mT2-tagged ASCL1 in many cells providing parallel evidence of cyclical fluorescence (Figures 4D and S5G, Supplemental Videos SV5 and SV6). Oscillations occurred with an average period of 202.5 ±9.9 min (mT2, period of highest probability 170 min) or 206.2 ±10.7 min (mNG, period of highest probability 200 min – Figure 4E). In replicating mouse NPs, *Ascl1* acts upstream of *Neurog2*,^53^ but human EEC precursors activate *ASCL1* far downstream of Neurog3 and we did not observe division of ASCL1^+^ fluorescent nuclei. A ∼12-h limit on robust live cell imaging precluded defining the number of oscillatory cycles; the large fraction of *ASCL1^+^*or *HES6^hi^* cells captured in every experiment (e.g., Figures 1F, 2E and 3A), coupled with a mean period of ∼3.5 h, suggests prolonged oscillation between *CREB3L1*-dominant and *NEUROD1^+^* phases (Figure 4F).

### Genomic binding of candidate pivotal TFs

Like ASCL1, NEUROD1 is present in NE cells of all tissues and in distinct sub-types of human NE cancers;^54–60^ as in mouse neurogenesis,^61,62^ *NEUROD1* appears late in the EEC trajectory. Alone or together with other TFs, ASCL1 converts mouse and human fibroblasts into neurons,^63,64^ in part by activating neuronal *cis*-elements within closed chromatin,^65^ while NEUROD1 promotes neural differentiation of embryonic stem cells^66^ and maturation of ASCL1-induced neurons.^67^ Both TFs display “pioneer” activity at large fractions of their binding sites in mouse or human neural cells differentiated *in vitro*^68–72^ and in human brain tumor cells.^73^ *Ascl1^-/-^* neonates lack stomach EECs (for unknown reasons – upstream *Neurog3* expression is preserved)^74^ whereas *Neurod1*-null mice have deficits only in intestinal S (secretin^+^) and I (cholecystokinin^+^) cells.^20^ In this light, our findings nominated ASCL1 and NEUROD1 as key successive TFs in human EEC differentiation, each possibly preparing chromatin for subsequent TF binding. We therefore used CUT&RUN (Ref. ^75^) to identify their binding sites in the genome 96 h after Tam exposure, when both TFs are present in sizable cell fractions (Figure S3D).

In two samples with ASCL1 antibody (Ab) and 4 independent replicates with NEUROD1 Ab, we identified 8,620 and 12,652 sites where one Ab or the other gave signals significantly stronger than IgG control. These numbers are on a par with those of binding sites detected by ChIP-seq in diverse cells^57,70,71^ and more than half the sites lie far from promoters (Figure S6A). Of the 15,429 unique binding sites, 18% preferred ASCL1 (A>N sites), 44.1% bound NEUROD1 more strongly (N>A), and 37.9% gave comparable signals with both Ab (A=N, Figure 5A). Shared sites are not surprising because E-box (CA**NN**TG) motifs for the two TFs differ by a single base: CA**G/CC**TG for ASCL1 (shared with bHLH factor TCF12) and CA**TC**TG for NEUROD1. Indeed, A>N and N>A sites were enriched for the respective motifs, while N=A sites, which included EEC-restricted loci *SYP* and *INSM1* (Figures 5A and S6B), were enriched for CA**T/CC**TG and other motifs (Figure S6C). Each k-mer was present at all three classes of binding sites, albeit in different proportions (Figure 5B). Micrococcal nuclease cutting patterns distinguish direct TF contact with DNA from indirect binding^76^ (in complex with some other sequence-specific TF – Figure 5C). By this measure, ASCL1 bound directly at its own optimal sequence and at one it shares with TCF12, but indirectly at the NEUROD1 motif, whereas NEUROD1 bound directly at both its own and the ASCL1 motif (Figure 5C, ETS and RFX motifs illustrate indirect binding). Thus, the two TFs occupy a large fraction of the same genomic sites, with NEUROD1 contacting DNA directly at most ASCL1 binding sites. Since transient *ASCL1* expression precedes *NEUROD1* and these TFs are co-expressed in only a tiny precursor fraction (Figure 1G), these findings imply that NEUROD1 replaces ASCL1 at most sites.

**Figure 5.**
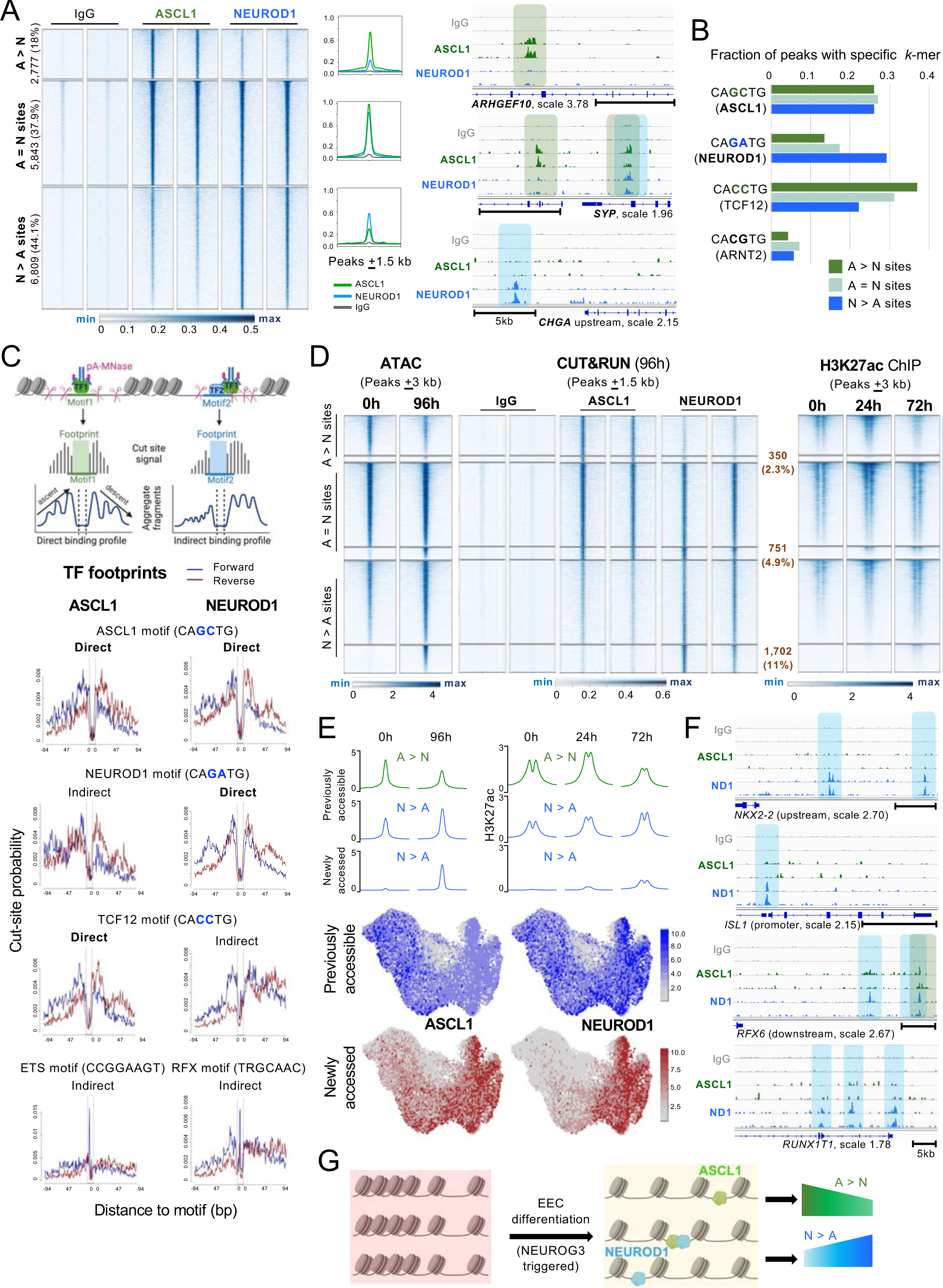
Genomic binding of ASCL1 and NEUROD1 and relation to accessible chromatin. (A) ASCL1 and NEUROD1 binding sites identified by CUT&RUN 96 h after Tam induction; 2 of 2 (ASCL1) or 4 (NEUROD1) replicates are shown and IgG provides a negative control. Sites preferentially bound by either TF are designated as A>N or N>A and sites occupied equally by both TFs as A=N. Aggregate plots show the average signals for ASCL1 and NEUROD1 at each class of binding sites and Integrated Genome Viewer (IGV) tracks at representative loci illustrate preferential or shared binding of ASCL1 (green) and NEUROD1 (blue). (B) Fractions of selectively bound and common regions that contain specific E-box (CANNTG) k-mer sequences within 120-bp windows around peak summits. (C) CUT&RUN-derived TF footprints. pA-MNase, bound to a TF antibody, cleaves DNA around binding sites differently when the TF contacts DNA directly at a motif of interest or indirectly in complex with some other TF. Lack of DNA sequence reads from a motif implies protection by the occupying TF, while flanking reads signal MNase accessibility. Direct TF binding at its motif gives equal numbers of reads on either side (equal slopes of ascent and descent), else binding is considered indirect. From the footprint patterns shown below, ASCL1 signals direct contact with ASCL1 and TCF12 motifs and indirect contact with the NEUROD1 (and control ETS-family) motifs, whereas NEUROD1 leaves a direct footprint at NEUROD1 and ASCL1 motifs but an indirect footprint at TCF12 (and control RFX-family) motifs. (D) ASCL1 and NEUROD1 binding sites (from CUT&RUN) clustered in relation to previously accessible and inaccessible regions, based on bulk ATAC-seq signals (Figure 2C) at the bound sites in untreated hISCs (0 h) or 96 h after initial Tam exposure. Only 2.3% to 11% of binding sites were previously inaccessible. Right, H3K27ac ChIP-seq data from the same binding sites in untreated cells and 24 h and 72h after induction of EEC differentiation. (E) Aggregate ATAC-seq (left) and H3K27ac ChIP seq (right) plots for previously accessible and newly accessed sites at the indicated times. snATAC signals at the two types of TF binding sites, projected onto the integrated UMAP from snATAC-seq (Figure 2D), confirm that sites appearing as “previously accessible” in bulk ATAC-seq had open chromatin before, whereas chromatin at “newly accessed” sites opened only after, *ASCL1* or *NEUROD1* expression. (F) IGV tracks showing NEUROD1, but little to no ASC1, binding at TF loci that are selectively enriched in the *NEUROD1^+^*and subsequent states, suggesting that *NKX2-2*, *ISL1*, *RFX6*, and other TF genes may be direct transcriptional targets of NEUROD1. (G) Model for selective and shared ASCL1 and NEUROD1 binding in largely pre-accessible regions and the consequences on subsequent chromatin accessibility. See also Figure S6.

If ASCL1 or NEUROD1 are pioneer TFs, the corresponding *cis*-elements would be inaccessible in resting hISCs and acquire access only after trace levels appear during EEC ontogeny.^77^ To test this possibility, we examined bulk ATAC- and ChIP-seq data (Figure 2D) from hISCs before and after NEUROG3 induction. Almost 93% of ASCL1-bound and 84.1% of NEUROD1-bound sites on day 4 were accessible in resting hISCs and the flanking nucleosomes carried H3K27ac (Figure 5D). Thus, although ASCL1 and NEUROD1 appear much after the onset of EEC differentiation, each TF binds mainly at pre-existing CRE; only a minority of bound sites open after they appear (Figure 5E). Subsequently, chromatin access and H3K27ac diminished at sites bound only by ASCL1; however, both access and H3K27ac increased at NEUROD1-bound sites (Figures 5D, 5E, and S6D), where new access was evident at loci that encode 8 of the 10 TF genes most enriched in the *NEUROD1^+^* cluster, but hardly any sites near (<100 kb) TF genes enriched in terminal EEC descendants (Figures 5F and S6B). Thus, ISCs are primed for EEC differentiation, ASCL1 acts transiently, and NEUROD1 binds TF loci associated with EEC maturation (Figure 5G).

### Precocious EEC differentiation in the absence of ASCL1 or HES6

We tested specific TF requirements in human EEC differentiation using CRISPR/Cas9 editing to disrupt *ASCL1* and, knowing that HES factors oscillate out of phase with ASCL1 in NPs,^53^ we also disrupted *HES6* (Figure S6E). Because EEC precursors activate *NEUROD1* downstream of *ASCL1* and the two TFs bind many of the same genomic sites, we also disrupted *NEUROD1* upstream of its C-terminal trans-activation domain^78^ (*NEUROD1^TAD^*) and, using different guides (gRNAs), its N-terminal DNA-binding domain (DBD). Amplicon sequencing confirmed gene targeting in >99% (*ASCL1* and *HES6*) and 85-87% (*NEUROD1*) of cells (Figure S6E). After NEUROG3 activation, pseudobulked RNA reads from snMultiomes confirmed absence of targeted transcript regions and immunoblots verified losses compared to control cells treated with scrambled gRNAs (Figure S6F).

Because *HES6* and *ASCL1* appear by the 3^rd^ day and both promote neuronal differentiation,^53^ we first examined *ASCL1*- and *HES6*-null cells on day 3, before *NEUROD1* is normally expressed. qRT-PCR analysis of both mutants showed 40- to >100-fold elevation of *NEUROD1*, *CHGA*, and non-EC peptide hormone mRNAs, which usually appear long after day 3 (Figure 6A – EC marker *TPH1* was not significantly altered). Immunoblotting confirmed a large increase in NEUROD1 and immunocytochemistry verified premature SST expression in *ASCL1*-null cells within 72 h of Tam exposure (Figure 6B and 6C). Mutant cell snMultiomes collectively showed premature expression of hundreds of mature EEC genes and intact expression of intermediate markers that ordinarily persist in mature EECs (examples in Figure 6D); these findings justified our integrated, nearest-neighbor consideration of control (scrambled gRNA) and mutant cells, with gene anchors transferred from normal EEC differentiation (Figure 1F). In this integrated analysis, transcriptomes of mutant cells classified as *NEUROD1^+^* or their mature descendants resembled their control counterparts but vastly outnumbered them (58% to 75% of mutant cells, compared to <25% of control cells, Figures S6G). Precocious *NEUROD1* expression and mature cell excess in both mutants thus indicate that, contrary to neurogenesis, HES6 and ASCL1 retard rather than promote EEC maturation, in part by restraining NEUROD1 activation.

**Figure 6.**
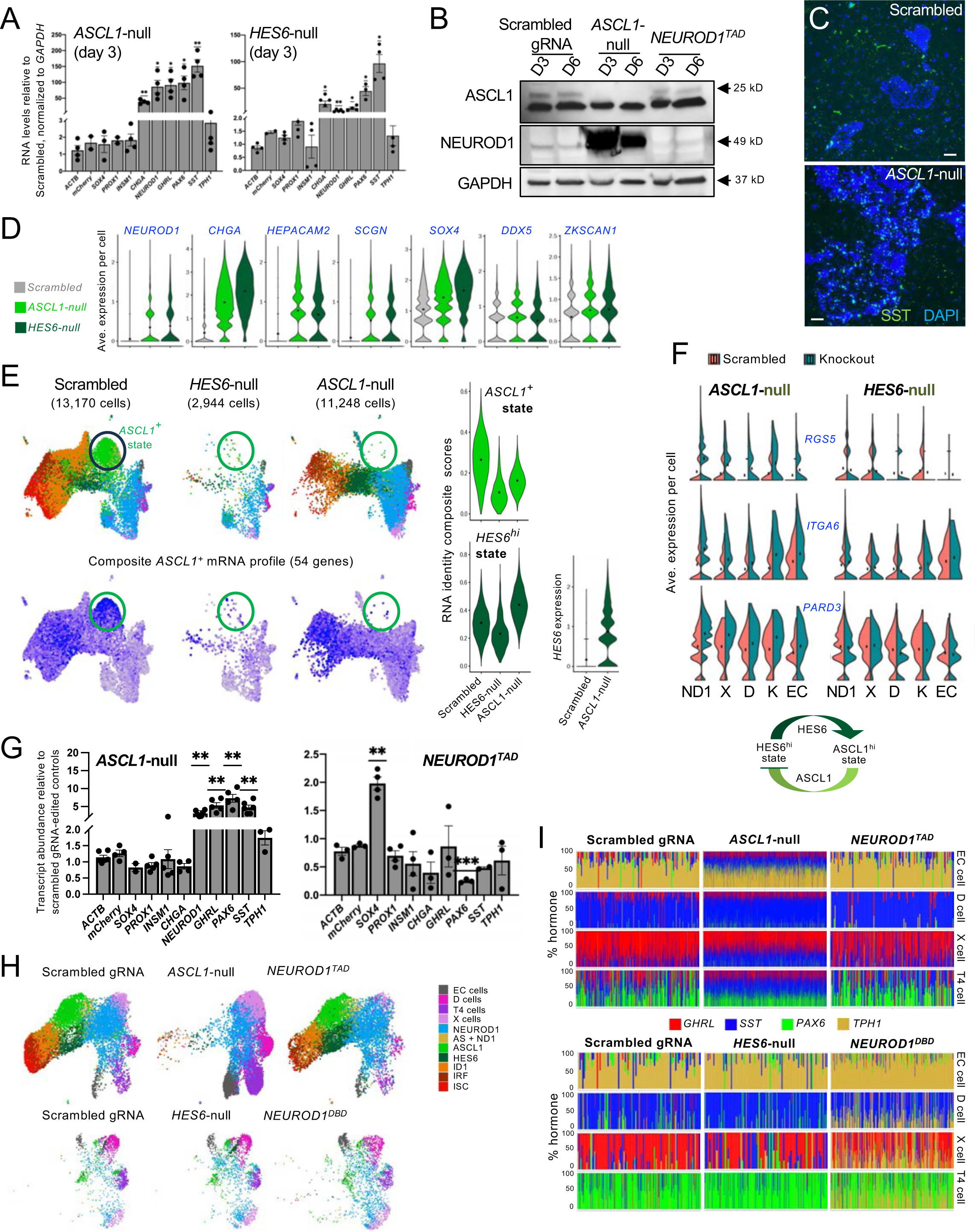
Precocious differentiation of *ASCL1*-null and *HES6*-null human EECs. (G) qRT-PCR analysis of edited *ASCL1*- and *HES6*-null cells relative to scrambled gRNA-edited controls (normalized to *GAPDH* levels) 72 h after NEUROG3 activation, showing a vast excess of terminal (*NEUROD1, CHGA*, *GHRL*, *PAX6* and *SST*) but not early (*SOX4*, *PROX1* and *INSM1*), EC (*TPH1*), or control (*ACTB* and *mCherry*) markers in both mutants. (means ±SEM, n ≥3 independent experiments, ** *p* <0.01, * *p* <0.05, t and Wilcoxon tests). (H) Immunoblot validation of TF gene editing (n=3 independent replicates). Protein lysates harvested 72 h and 144 h after Tam exposure from CRISPR/Cas9-edited cells to disrupt *ASCL1* or the trans-activation domain (TAD) of *NEUROD1* and from control (scrambled gRNA) cells were blotted with ASCL1, NEUROD1 or GAPDH (loading control) Ab. Expected M_r_ for each protein is indicated. In *ASCL1*-null cells, ASCL1 is eliminated and NEUROD1 is massively increased. In *NEUROD1^TAD^* cells, validated by amplicon sequencing (Figure S6E), the protein product of gene targeting has the same M_r_ as the wild-type protein. (I) Immunofluorescence showing precocious and widespread SST expression in *ASCL1*-null cells 72 h after NEUROG3 activation (n=3 independent experiments, scale bar 100 µm). (J) scRNA analysis of *ASCL1*-null and *HES6*-null cells relative to scrambled gRNA-edited controls (normalized to *GAPDH* levels) 72 h after NEUROG3 activation, showing massive increase in both mutants of terminal (*NEUROD1*, *CHGA*, *HEPACAM2*, *SCGN*) compared to early or broadly expressed (*SOX4*, *DDX5*, *ZKSCAN1*) markers. (K) Merged UMAP of snMultiome-seq derived cell clusters 72 h after Tam exposure in *HES6*-null, *ASCL1*-null, and control cells. Below, expression of the composite 54-gene panel enriched (Table S3, not all genes are exclusive) in the *ASCL1^+^* state is projected onto each UMAP. Right, levels of the 54-gene *ASCL1^+^* and 43-gene *HES6^hi^* signatures are quantified. (L) Expression of representative *HES6^hi^* signature genes in *ASCL1*-null and *HES6*-null maturing *NEUROD1^+^* (ND1) cells and in their mature descendants. The sum of our findings indicates that high HES6 levels activate *ASCL1* expression, which in turn represses the transitional *HES6^hi^* state, generating the observed oscillation. (M) qRT-PCR analysis of edited *ASCL1*- and *NEUROD1^TAD^* cells relative to scrambled gRNA-edited controls (normalized to *GAPDH* levels) at 144 h, showing an excess of terminal (*NEUROD1, GHRL*, *PAX6* and *SST*) but not early (*SOX4*, *PROX1* and *INSM1*) markers in *ASCL1*-null but not in *NEUROD1^TAD^* cells. (means ±SEM, n ≥3 independent experiments, ** *p* <0.01, calculated from t and Wilcoxon tests). (N) Merged UMAP of snMultiome derived cell clusters 144 h after NEUROG3 induction in each mutant and in control (top n=10,707, bottom n=1,914) cells. Clusters are labeled according to gene anchors transferred from scRNA-seq of wild-type EEC differentiation (Figure 1). (O) Relative expression of individual hormone RNAs in clusters meeting criteria to be classified as EC, D, X or T4 cells. Each column within a block represents relative hormone expression in one cell. *SST* is expressed ectopically in multiple *ASCL1*-null EEC types. *NEUROD1^TAD^* shows only minor reductions in *GHRL* expression in X cells; however, in *NEUROD1^DBD^* cells, ectopic *TPH1* mRNA, which signifies EC differentiation, is present in each non-EC type. See also Figure S6.

Although nearest-neighbor analysis classified some *HES6*- and *ASCL1*-mutant cells as *ASCL1^+^* precursors (Figure S6), the corresponding sector of the integrated UMAP was notably bare (Figure 6E). Moreover, the 54-gene *ASCL1^+^* state signature (Figure 3F) was markedly attenuated (Figure 6E – as some of these genes are enriched but not restricted to *ASCL1^+^* cells, the collective signal extends beyond the *ASCL1^+^* cluster, and preservation of that signal points to selective loss of the *ASCL1^+^*precursor state). Conversely, the 43-gene *HES6^hi^* transcriptional signature, which is less specific than its *ASCL1^+^* counterpart (Figure 3F), was mildly diminished in *HES6*-null but amplified in *ASCL1*-null derivatives, where *HES6* expression was elevated in all cell fractions (Figures 6F and S6H) and 3 times as many cells classified as *HES6^hi^*than in the control (Figure S6G). Some *HES6*-synexpressed genes, e.g., *RGS5*, were suppressed in *HES6*-deficient EECs, while others (e.g., *ITGA6* and *PARD3*) were not (Figure 6E). Together, these findings directly implicate ASCL1 in driving its namesake state and HES6 (which lacks a DBD but counters gene repression by other Hes-family TFs such as HES1)^51^ in promoting that state. Conversely, ASCL1 normally represses *HES6* expression (Figure 6E) and that reciprocity likely underlies its oscillation.

### Compromised EEC fidelity in the absence of specific TFs

Not only did *HES6*- and *ASCL1*-null EECs mature at least 3 days early, but X and D cell features were represented unfaithfully (Figure S7A). To extend those observations and because *NEUROD1* appears by day 4, we examined consequences of TF deficiencies 6 days after Tam induction. qRT-PCR analysis of Tam-treated *ASCL1-*mutant cells showed 5- to >10-fold elevation of terminal EEC transcripts, consistent with precocious maturity, while *NEUROD1^TAD^-*derived cells showed small changes (Figure 6G). For sn studies, we paired *ASCL1*-null and *NEUROD1^TAD^*cells with a control (scrambled gRNA) in one experiment, where EEC differentiation was complete but slow in all samples, and *HES6*-null and *NEUROD1^DBD^*cells with a control in another experiment, where EECs differentiated at the usual speed, yielding mainly terminal cells on day 6. Individual cell clusters in both controls resembled each other and untreated hISC^NEUROG^^3^ (Figure S7B), allowing mutant lines to be compared fairly against one another.

Nearest-neighbor analysis of mutant snMultiomes, each integrated with its own control, identified cells corresponding to each interim and terminal EEC state (Figure 6H). As expected from precocious EEC maturation, precursor cells that lingered in control and *NEUROD1^TAD^* were absent from *ASCL1*-null derivatives, where the 4 terminal EEC types dominated (>55% compared to <20% in controls, Figure 6H). Mutant cells tagged as EC, D, X or T4 showed highly anomalous hormonal repertoires. *ASCL1*-deficient EC cells aberrantly expressed peptide hormone genes, while X cells expressed high *SST* levels (Figure 6D); *HES6*-deficient X cells also expressed ectopic *SST* at the expense of *GHRL*; and *NEUROD1^DBD^* mutant non-EC cells broadly expressed the EC marker *TPH1* (Figure 6I). These anomalies do not reflect computational misallocation of cell types because those combinations of high hormone transcripts are rare in any normal population (Figure 1) or scrambled controls. Indeed, Ab staining revealed SST peptide in abnormally high fractions of *ASCL1*- (*P* <0.001) and *HES6*- (*P* <0.0001) null cells at the expense of GHRL and an elevated fraction of SST- and TPH1-co-expressing cells in *NEUROD1^DBD^*mutants (*P* <0.01, Figure 7A). Moreover, *ASCL1*- null cells showed premature expression of terminal EEC transcripts in cells with the features of *NEUROD1^+^* precursors and *NEUROD1^DBD^* mutants expressed the EC marker *TPH1* broadly (Figure 7B). Ectopic expression in mutant EECs was not confined to hormone genes: the mRNA signatures of all terminal non-EC (Figure S2G), especially X and D cells, were evident in all *ASCL1*-null derivatives, including *NEUROD1^+^*precursors and EC cells (Figure 7C). Conversely, the 532-gene X-cell fingerprint, which had diminished in *HES6*-null EECs by 3 days (Figure S6H), was markedly attenuated on day 6 (Figure 7C); the 756-gene EC signature extended into *NEUROD1^TAD^*and even more appreciably into *NEUROD1^DBD^*non-EC cells (Figure 7D). Thus, beyond restraining terminal EEC differentiation, ASCL1, HES6 and NEUROD1 ensure accurate expression of terminal genes normally restricted to single EEC types. Abnormalities were most pronounced upon ASCL1 loss, with ASCL1 and NEUROD1 deficiency exerting reciprocal effects on EC versus non-EC, and ASCL1 or HES6 deficiency having reciprocal effects on X versus D cell, character. Because ASCL1 loss perpetuates the *HES6^hi^* state and activates *NEUROD1* prematurely (Figures 6E and 6F), its oscillatory expression likely mediates both delayed and faithful EEC lineage allocation.

**Figure 7.**
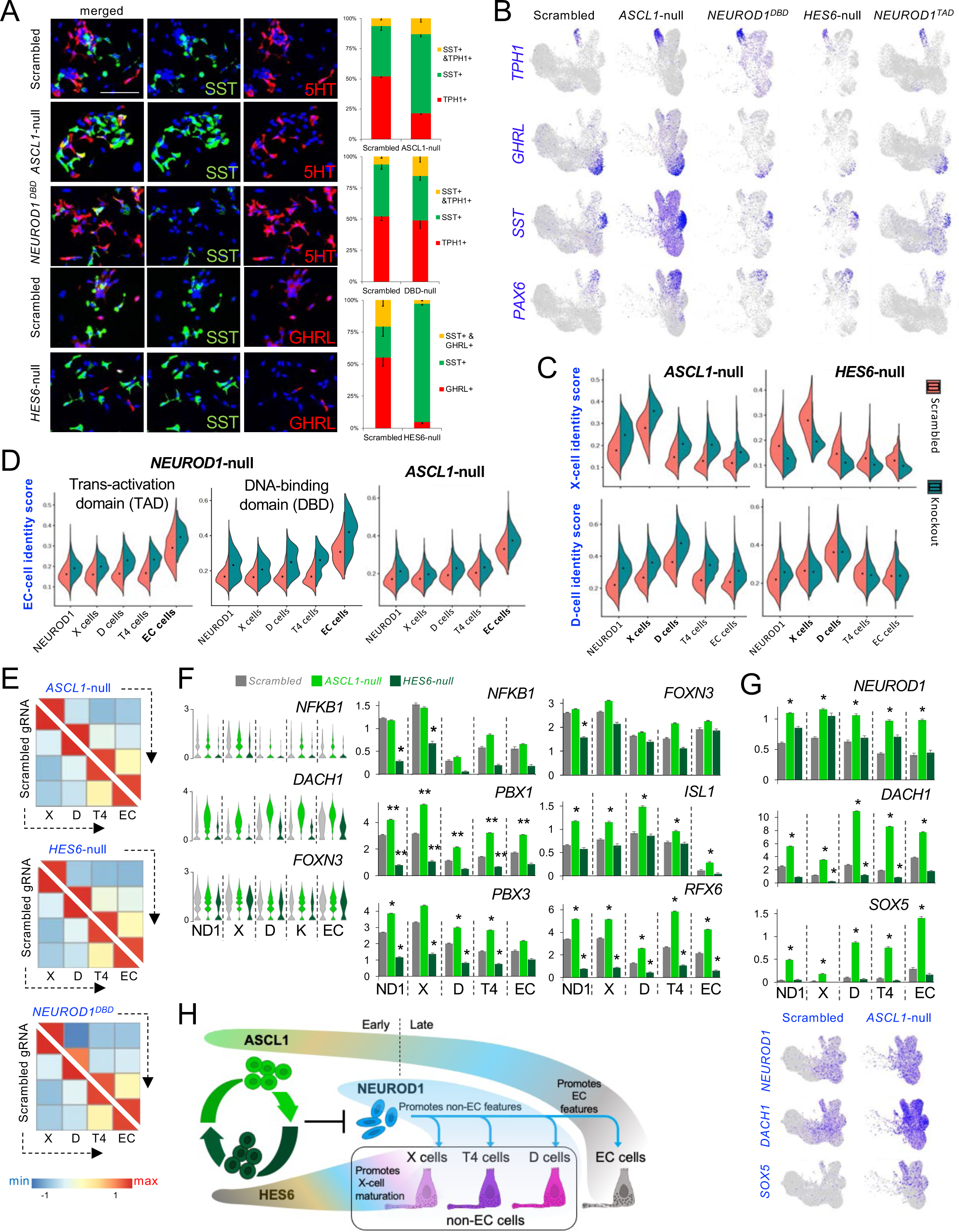
EEC lineage infidelity in the absence of specific TFs. (A) Co-immunofluorescence for Somatostatin (SST) and 5-hydroxytryptamine (5HT) in *ASCL1*- null, NEUROD1^DBD^, and control (scrambled gRNA) cells and for SST and Ghrelin (GHRL) in *HES6*-null and control cells. 50 µm scale bar applies to all images. Cell proportions stained with one or both markers 144 h after Tam exposure are quantified, showing increased SST^+^ cell fractions in the absence of ASCL1; increased SST^+^ TPH1^+^ (double-positive) fractions in the absence of ASCL1 or the NEUROD1 DBD; and a marked drop in GHRL^+^ cells in the absence of HES6. (% mean ±SEM, n: at least 3 independent experiments, with 20-100 cells counted per sample). (B) Expression levels of EC marker *TPH1*, X-cell marker *GHRL*, D-cell marker *SST*, and T4 cell marker *PAX6* mRNAs in control cells and each mutant, projected onto an integrated UMAP. The data show aberrantly broad expression of *TPH1* in *NEUROD1^DBD^* derivatives, of *GHRL* and *SST* in *ASCL1*-null derivatives, and reduced *GHRL* expression in *HES6*-null derivatives. (C) Transcriptional identity scores of individual terminal EEC types (from Figure S2G) in mutant derivatives. X-cell identity extends into *ASCL1*-null D and T4 cells and is substantially diminished in *HES6*-null X cells. *ASCL1*-null, but not *HES6*-null, derivatives show elevated expression of the D-cell signature. (D) EC identity extends into non-EC derivatives in *NEUROD1*-mutants (subtly upon disruption of the TAD alone, notably upon disruption of the DBD) but is unperturbed in *ASCL1*-null EECs. (E) Comparisons of accessible X, D, T4, and EC enhancers in EECs derived from *ASCL1*-null, *HES6*-null, and *NEUROD1^DBD^* cells. Perturbation of open chromatin is negligible. (F) Expression of many TF genes differentially expressed in normal EECs (Figure 2B) is disturbed, often reciprocally, in *ASCL1*- and *HES6*-null EECs. These changes are shown in bar graphs and, for selected examples, also in violin plots. Pan-EEC marker *FOXN3* is a representative control (unperturbed) gene. * |log_2_ fold-change| >0.25 and *p* <0.001 using Wilcoxon rank sum test compared to scrambled gRNA-edited controls. n.s., not significant. (G) Substantially increased expression of *DACH1* and *SOX5* in derivatives of both mutants and of NEUROD1 more prominently in *ASCL1*- than in *HES6*-null EEC derivatives. In the feature plots, expression levels of these TF genes are projected on the integrated UMAP. (H) Summary of distinct ASCL1, HES6, and NEUROD1 functions early and late in human EEC differentiation. See also Figure S7.

### Chromatin accessibility and downstream TFs in mutant EECs

As mature EECs share many CREs and TFs (Figures 2B, 2E and 4C), the anomalies resulting from TF losses could reflect extensive or limited chromatin aberrations. Even in TF-deficient EECs that mis-expressed 150 to 650 genes (log_2_ fold-change >0.3, *P*_adj_ <0.05), snMultiomes revealed negligible defects in global profiles of open chromatin (Figure 7E) and few CREs showed aberrant chromatin access (log_2_ fold-change >0.2, *P*_adj_ <0.05, Figure S7C). Even the minority of ASCL1- (7%) or NEUROD1-bound (∼16%) sites that became accessible after those TFs are expressed (Figure 5E) remained open in the absence of either factor (Figure S7D), indicating absence of non-redundant pioneer activity. For additional directed interrogation of CREs, we linked the TSSs of all mis-expressed genes, including ectopic hormone products, to accessible sites within 500 kb and derived regulatory module scores, which were unchanged in mutant cells (Figure S7E). Thus, ASCL1, HES6, and NEUROD1 losses profoundly affect EEC phenotypes without appreciably disturbing chromatin accessibility, a finding compatible with relatively uniform chromatin landscapes in maturing *NEUROD1^+^*cells (Figure 2F).

Instead, we detected considerable aberration among the 38 TF transcripts normally present at sufficient levels in selected EEC types, plus others expressed at low basal levels (see Figure S7C). For example, X cell-enriched *NFKB1*, *HES4*, *EPAS1*, *PBX1*, and *PBX3* transcripts were reduced in *HES6*-null X cells and most were reciprocally increased in *ASCL1*-null cells (Figures 7F and S7F). These reductions do not simply signify defective X-cell differentiation because some of the TFs are normally also expressed in other EECs (Figure 2B), which were little affected upon HES6 loss, and their expression was reduced in all *HES6*-null EECs, including *NEUROD1^+^* precursors; *FOXN3*, which is expressed broadly, represents the many TF genes that are affected minimally or not at all. Conversely, D cell-enriched *ISL1* and *MAFB* and T4 cell-enriched *RFX6* were elevated in all *ASCL1*-null EECs, including EC cells, without or with (respectively) reciprocal diminution in *HES6*-null derivatives (Figures 7F and S7F). Notably, the few TFs strictly confined to one EEC type, such as *HHEX* or *POU2F2* (D cells), *FEV* or *MNXI* (EC cells), and *PAX6* or *PDX1* (T4 cells) were largely unaffected (Figure S7G). Rather, the most perturbed TF genes (*PBX1*, *PBX3*, *RFX6*, *MAFB*) normally appear in more than one EEC type (Figure 2B) and, in keeping with the preserved global profiles of open chromatin, discernible changes in local chromatin accessibility did not accompany their perturbation (Figure S7H). Two low-expressed TF genes not enriched in any normal cluster, *DACH1* and *SOX5*, were substantially increased in *ASCL1*-null derivatives (Figure 7G). As the sparsity of RNA reads in any single cell makes it difficult to appreciate the scale of TF transcript changes, it is instructive to compare *DACH1* and *SOX5* with *NEUROD1*, for which bulk cultures revealed large increases on day 3 in both *ASCL1* and *HES6* mutants (Figure 6).

The scale and reciprocal patterns of TF gene dysregulation hence mirror, and likely underlie, the reciprocal differentiation defects in mutant cells, related in turn to precursor state oscillations (Figure 7H). The sum of our findings suggests that by delaying *NEUROD1* activation, ASCL1 and HES6 prepare EEC precursors to express TFs in the proper cellular domains that drive distinctive terminal transcriptomes.

## DISCUSSION

The present understanding of EEC ontogeny rests heavily on observations made in mice, where cell types and regulatory networks differ from human EECs,^29^ and on investigation of relatively few cells differentiated *in vitro*. We examined mRNA and chromatin trajectories in substantially at least 2 orders of magnitude more cells than studied in the past, including abundant precursors not previously captured along the continuum of human ileal EEC differentiation. We thus identify the basis for efficient NEUROG3-driven differentiation of human ISCs into diverse mature EECs that express unique transcriptional programs and the means to synthesize serotonin or distinct hormones. The earliest events encompass attenuation of the ISC program, including cell replication. EEC-restricted enhancers and genes are activated only after much of that program is abrogated and after a period of ASCL1 oscillation associated with limited chromatin and transcriptional flux. Distinct EEC types subsequently mature from *NEUROD1^+^* precursors in which a majority of CREs for EEC genes, previously inaccessible to TFs, have become accessible. Although smaller numbers of CREs acquire selective access in individual terminal EEC types, the lineage diversifies from a shared platform of open chromatin present in *NEUROD1^+^* cells, through the actions of ∼40 TF genes that appear in overlapping combinations in descendant EEC types. This common CRE platform explains the ready *in vivo* conversion from production of one major hormone to another;^16^ it also explains why TF-deficient EECs in our study had extensively disrupted RNA profiles with little disturbance in open chromatin. Overlapping TF expression domains, coupled with distinct perturbations when we disrupted antecedent cell states, support the idea that specific TF combinations underlie terminal EEC differentiation. Although human and murine EECs differ in many respects, other elements are conserved,^16^ and EEC findings in scattered mouse TF gene knockouts^20–24^ are compatible with TF combinations expressed in human *NEUROD1^+^* descendants. hISC^NEUROG3^ cells provide a facile model to investigate additional TFs and candidate EEC determinants.

Although EEC types are recognized by their dominant hormone product, they have long been known to often express more than one hormone,^36,37^ as confirmed recently at single-cell RNA resolution.^16,26^ However, many hormone pairs have opposing effects, for example on appetite, exocrine pancreas stimulation, or glycemia.^8,14^ It is therefore important that any EEC tailor its hormonal repertoire to match its environmental sensors and secretory capacity; moreover, *TPH1* and other transcripts are exquisitely restricted to EC, which are unambiguously distinct from non-EC, cells. Compared to cells treated with a non-specific gRNA, our findings implicate *ASCL1*, *HES6* and *NEUROD1* in EEC fidelity, with ASCL1 and NEUROD1 losses showing opposite effects on EC vs. non-EC character and HES6 and ASCL1 losses showing opposite effects on X- vs. D-cell character. The latter reciprocity is noteworthy because *ASCL1^+^* precursors and ASCL1 expression oscillate with respect to *HES6^hi^* cells, loss of either factor markedly expedited EEC maturation at the expense of fidelity, and nearly every TF transcript reduced in *HES6*-null EEC derivatives was increased in *ASCL1*-null derivatives. TFs expressed downstream of ASCL1 appear in specific combinations in individual EEC types, and our findings suggest that ASCL1 oscillation before the advent of *NEUROD1^+^* precursors provides a quality control delay, ensuring that downstream TFs distribute properly and craft terminal EEC identities accurately. ASCL1 oscillation thus serves a different purpose in EECs than in NPs, where it promotes cell replication at the expense of differentiation.^53^ Of note, the focus of our study on CRE and transcriptional dynamics did not consider additional post-translational modulation of TFs. Differential TF, especially bHLH protein, phosphorylation influences differentiation of diverse cell types,^53,79^ likely including adult human EECs.

The moniker ‘neuro-endocrine’ (NE) derives from defining attributes that EECs share with pancreatic islet cells (metabolic homeostasis, response to food, related peptide hormones) and neurons (active environmental sensing, electrical excitability, packaging of products in vesicles that fuse with cell membranes after specific stimuli). Primitive neurogenic modules in peptidergic cells found in non-bilaterian placozoans^80^ may be the evolutionary forebears of neurons as well as gut, pancreas, and lung endocrine cells. Our findings highlight that, despite the related outlines and similar TF usage in mouse and human EECs and neurogenesis, there are notable tissue and species differences. Relations among NEUROG3, ASCL1, NEUROD1, and their homologs indicate that specific functions and epistatic relations need not extrapolate from one tissue to another or from embryos to adults. *Neurog3* triggers both mouse pancreatic islet cell development (human beta cells are less dependent^81^) and intestinal EEC specification, but whereas *Neurod1* acts immediately downstream in the fetal pancreas,^20^ it acts far downstream in adult human EECs, where we find that *ASCL1* restrains *NEUROD1* activation. While NEUROG3, and especially NEUROD1, pioneer chromatin at mouse and human neuronal CREs,^68,70–72^ they pioneer only a small fraction of all binding sites in human EECs, binding principally at enhancers that were accessible in ISCs lacking either TF. Additionally, absence of ASCL1 arrests neurogenesis, while blocking its oscillation drives proliferating NPs to exit the cell cycle, activate *Neurog3* homologs *Neurog1/2*, and differentiate into neurons.^53^ In human EECs by contrast, ASCL1, acting downstream of NEUROG3, oscillates in *post-mitotic* precursors and its loss *accelerates* NEUROD1 expression and EEC differentiation. Knowledge of these distinctions will assist in engineering cellular EEC therapies and as a framework to approach human intestinal, pancreatic, and other NE cancers, which commonly express ASCL1 or NEUROD1 to the exclusion of the other, with different prognostic and mechanistic implications.^57–60^

## Author contributions

P.N.P.S., Q.Z. and R.A.S. conceived and designed the study; P.N.P.S. and W.G. characterized EEC differentiation; P.N.P.S. performed most experiments, some with help from S.B.; S.M. and R.H. assisted with computational analyses; A.W.L and C.A.M performed MIRA analyses; P.C., M.G., M.B. and H.W.L. assisted with scATAC or multi-ome studies; M.O. assisted with TF gene editing; P.N.P.S. and R.A.S. interpreted data and wrote the manuscript, which all authors edited.

## Acknowledgments

Supported by a Peterson Accelerator grant (R.A.S. and Q.Z.) from the Neuroendocrine Tumor Research Foundation and by NIH grant R01 DK082889 (R.A.S.). We thank L. Perrault for help with immunocytochemistry; K. Huang, P. Speliakos, N. Serbyn, D. Pellman, J. Heneghan, and J. Turner for guidance on live cell imaging; S. Henikoff for the gift of MNase-conjugated Protein A; and staff in the organoid core of the Harvard Digestive Diseases Center (P50 DK034854) and the flow cytometry core at the Dana-Farber Cancer Institute.

## MATERIALS AND METHODS

### Lead Contact

Further information and requests for resources and reagents should be directed to the Lead Contact, Ramesh A. Shivdasani.

### Materials availability

Plasmids to express Neurog3 (Ref.^102^) were purchased from Addgene. Human organoid lines generated in this study are available and can be requested from the Lead Contact; a Materials Transfer Agreement may be required.

### Data and code availability

Sequencing data analyzed in this paper have been deposited in the Gene Expression Omnibus (GEO) repository under the accession code GSE238276.

## EXPERIMENT MODEL AND SUBJECT DETAILS

### Human biopsy samplesC

Normal human duodenum and ileum biopsy tissue samples were obtained from donors who provided informed consent, under protocols approved by ethics committees at Boston Children’s Hospital or the Dana-Farber/Harvard Cancer Center. The study complies with pertinent ethical regulations concerning research use of de-identified human tissues.

## METHOD DETAILS

### Generation and culture of primary 2D hISC lines

The epithelium was manually stripped from human intestinal biopsy samples collected in cold phosphate-buffered saline (PBS), minced, and digested with 200 IU/ml collagenase type I (Worthington, LS004196) in Hank’s Balanced Salt Solution (HBSS) at 37°C for 30-60 min with agitation and vigorous pipetting. Dissociated tissues were filtered through a 100 µm filter and centrifuged at 500 *g* for 5 min. Pelleted cells were washed with F12K medium and cultured as 3D organoids in Matrigel (Corning, 356231). When organoid growth had slowed after a few passages, cells were pelleted, washed with F12K media, resuspended in hISC culture medium, (90% basal media - containing 60% Dulbecco’s Modified Eagle Medium (DMEM, high glucose, Thermo Fisher Scientific, 11965092), 20% F12K (Thermo Fisher Scientific, 21-127-022), and 20% fetal bovine serum (FBS, Corning 35-010-CV); supplemented with 10% Rspo2 conditioned medium, 10 mM nicotinamide (Sigma, N5535), 25 µM Primocin (Invivogen, ant-pm-1), 1 µM A8301 (Sigma, SML0788), 5 µg/ml Insulin (Sigma I0516), 10 µM Y27632 (LC Laboratories Y5301), 1µM DMH1 (Sigma D8946), 50 ng/ml EGF (R&D, 236EG200) and 2 µM T3 (Sigma, T3697)), seeded on dishes coated with mitomycin C (Cayman, 11435) inactivated DR4 mouse embryonic fibroblasts (MEFs, American Type Culture Collection, SCRC-1045), and expanded and maintained as 2D colonies at 37°C in a 7.5% CO_2_ incubator. Culture medium was replaced every 2-3 days. Primary 2D hISC colonies were passaged every 4-6 days by incubating for 10-12 min with gentle pipetting in TrypLE (Life Technologies, 12604021), neutralized with 10% FBS in Dulbecco’s Modified Eagle Medium (DMEM, Life Technologies, 11965092), and centrifuged at 300 *g* for 5 min. Pelleted cells were resuspended in hISC culture medium, split at 1:3 ratio, and seeded on dishes coated with mitotically inactive MEFs for continued expansion.

### Generation of NEUROG3 inducible hISCs

*Neurog3* was amplified from mouse genomic DNA and cloned into the lentiviral vectors pLenti-TetO-Neurog3 and pLenti-EF1α-Neurog3ER-Puro^R^-mCherry. Lentiviral particles (10^8^ transduction units/ml) were produced using HEK293FT cells, as described previously.^102^ We used two independent hISC organoid lines to generate tamoxifen- or doxycycline-inducible Neurog3 lines. We transduced 2D hISC cultures with lentiviral particles carrying the polycistronic cassette TetO-Neurog3 and PGK-rtTA-Blast^R^ or EF1α-Neurog3ER-Puro^R^-mCherry in the presence of 10 µg/ml polybrene, as described previously.^102^ Transduced colonies were selected 48 h later using blasticidin (10 µg/ml) or puromycin (2-4 µg/ml) and stably transduced cells were visualized by constitutive mCherry expression from the polycistronic cassette EF1α-Neurog3ER-Puro^R^-mCherry. Engineered hISCs were pulsed for 48 h with Doxycycline (1 µg/ml) to induce TetO-driven *NEUROG3* transcription or with 4-hydroxytamoxifen (1 µM, Sigma-Aldrich, H7904) to activate NEUROG3.

### Fluorescence-activated cell sorting

mCherry^+^ and DAPI^-^ cells were sorted using a BD FACS M Aria II instrument with 70 µm nozzle. Cells were collected in FACS buffer (0.9% Glucose, 10 mM HEPES, 10 µM Y27632, 2% FBS, and N-acetyl-L-cysteine in HBSS) and washed with PBS before further analyses.

### RNA extraction, quantitative PCR, bulk-RNA and single-cell RNA sequencing

We sorted 2 x10^5^ DAPI^-^ mCherry^+^ cells from the specified time points, washed the cells in PBS, and extracted total RNA using the Qiagen RNeasy Micro Kit (74004). For quantitative RT-PCR, complementary DNA was generated using the SuperScript^TM^ III First-Strand Synthesis System (Life Technologies, 18080051) and quantified using Power SYBR Green Kit (Life Technologies, 4367659). Primers used for cDNA quantification are listed in Table S6. RNA libraries for bulk-RNA sequencing were generated using Novogene services and sequenced on an Illumina HiSeqX instrument to generate paired-end 150-bp reads. For scRNA-seq, induction of NEUROG3 in cells from different times was staggered to allow harvesting and processing on the same day. ∼10^4^ DAPI^-^ mCherry^+^ cells were loaded on the 10x Genomics chip in separate wells for each time point analyzed and processed through Chromium Controller to generate single-cell gel bead emulsions. Libraries for single-cell RNA were generated as described by the 10x Genomics protocol, Chromium Next GEM Single Cell 3’ v3.1 (PN-1000128) and sequenced on an Illumina NovaSeq instrument (Novogene).

### Immunocytochemistry

Cells were seeded on coverslips coated with inactive MEFs and placed in 12-well dishes. At the desired time, cells attached on coverslips were washed with PBS, fixed in 4% paraformaldehyde for 10 min at room temperature, washed again with PBS, blocked using 5% donkey serum in PBS for 1 h at room temperature, and then incubated with CHGA (Abcam, ab15160, 1:200), SST (Dako, A0566, 1:500), 5-HT (Immunostar, 20080, 1:500) or GIP (Santa Cruz, sc-23554, 1:100) antibodies in blocking buffer (5% donkey serum in PBS) overnight at 4°C. Cells were washed 3 times with PBS containing 0.1% triton (PBST) and incubated with Alexa-Fluor 488-conjugated secondary antibodies in blocking buffer for 2 h at room temperature. Cells were washed, nuclei were stained with DAPI, coverslips were mounted on slides in mounting medium (Vector Laboratories, H-1000-10), and representative images were captured using a Nikon Widefield Ti2 Eclipse microscope. Images were processed using ImageJ^92^.

### Immunoblotting

Cells were lysed in RIPA buffer with Halt protease and phosphatase inhibitor (Thermo Scientific, 1862495, 1862209) for 30 min on ice. Lysates were pelleted at 16,000 *g* for 20 min at 4°C and the supernatant was collected and stored at −20°C. Protein extracts were quantified using BCA Pierce Kit. 20 µg of protein was boiled in 4X Laemmli buffer (Bio-Rad) for 5 min at 95°C, centrifuged for 1 min at 16,000 *g*, loaded onto 5-12% gradient gels (BioRad), and separated by electrophoresis at 100-150V for 45-60 min in 1X Tris-Glycine-SDS Running buffer (Boston BioProducts). Proteins were transferred to methanol treated PVDF membranes (Bio-Rad) at 70 V for 90 min at 4°C in 1X Transfer buffer with 20% methanol (Boston BioProducts). PVDF membrane were then blocked using 5% milk in TBST, incubated with primary antibodies overnight at 4°C, washed, and then incubated with horseradish peroxidase-conjugated anti-rabbit or anti-mouse IgG. Protein bands were visualized using enhanced chemiluminescent substrate (GE healthcare) and imaged on a LS Quant 4000 instrument (GE healthcare). ASCL1 antibody (Ab) was from Abcam (ab74065; 1:500); all other Ab were from Cell Signaling Technology: NEUROD1 (4373, 1:500), pSTAT1 (7649T, 1:1000), pSTAT2 (4441T, 1:1000), pSTAT3 (9145, 1:2000), STAT1 (14994T1, 1:1000), STAT2 (72604, 1:2000), and STAT3 (4904, 1:2000). Controls included GAPDH (Santa Cruz, sc25778, 1:1000), ACTB (Sigma, A5441, 1:1000) and normal rabbit IgG (Millipore, 12-370).

### Bulk and single-cell ATAC-sequencing and single-cell multiome ATAC + gene expression

Bulk ATAC analysis was performed using 5 x10^4^ DAPI^-^ mCherry^+^ cells sorted at the specified times and washed in PBS, followed by the Omni-ATAC protocol.^103^ Briefly, cells were lysed in 50 µl ice-cold ATAC-resuspension Buffer (RSB, contains 10 mM Tris-HCl ph7.4, 10mM NaCl, 3mM MgCl_2_, 0.1% IGEPAL, 0.1% Tween-20 and 0.01% digitonin) for 3 min on ice. Cells were then washed in ATAC-RSB buffer without IGEPAL and digitonin. Nuclei were pelleted by centrifugation at 500 *g* for 10 min at 4°C, resuspended in 50 µl of transposition mix (25 µl TD buffer, 2.5 µl 2µM TDE1 Tagment DNA enzyme (Illumina, 20034197), 16.5 µl PBS, 0.5µl 1% digitonin, 0.5 µl 10% Tween and 5µl nuclease free water) and incubated at 37°C for 30 min, followed by DNA purification using the MinElute PCR Purification Kit (Qiagen, 28004). Libraries were amplified as described,^40^ quantified by qubit dsDNA high-sensitivity assay (Life Technologies), assessed using High Sensitivity DNA Kit (Agilent) on a Bioanalyzer 2100 instrument (Agilent), and sequenced on a Illumina HiSeqX instrument (Novogene) to generate paired-end 150-bp reads. For single-cell ATAC and single-cell Multiome ATAC + Gene Expression sequencing, 2 x10^5^ DAPI^-^ mCherry^+^ cells were sorted, and intact nuclei were extracted as described above for bulk-ATAC. Induction of NEUROG3 in cells from different times was staggered to allow harvesting and processing on the same day. ∼10^4^ nuclei were loaded on the 10x Genomics chip for each time point. Libraries for single-cell ATAC and single-cell Multiome were generated as described in protocols PN1000176 and PN1000285, respectively (10x Genomics).

### ChIP-seq

ChIP-seq was performed on cross-linked chromatin as described previously.^104^ Briefly, ∼7.5 x10^5^ sorted DAPI^-^ mCherry^+^ cells were cross-linked at room temperature in 1% formaldehyde (Sigma, F8775) for 15 min, lysed in ChIP sonication buffer (50 mM Tris-HCl, 0.1% SDS, 10 mM EDTA and 1x Roche EDTA-free protease inhibitor) and sonicated in a Covaris E210 sonicator at 4°C for 50 min with 5 min on/off cycles to obtain 200-800bp DNA fragments, and incubated overnight with H3K27ac antibody (Active Motif, 39135) at 4°C. Antibody-bound DNA chromatin complex was incubated with a mixture of 15 µl Protein A and 15 µl Protein G magnetic beads (ThermoFisher, 10002D and 10004D) for 4h at 4°C, washed sequentially twice in low-salt buffer (20 mM Tris-HCl pH 8.1, 2 mM EDTA, 0.15 M NaCl, 0.1% SDS, 1% Triton X-100), once in high-salt buffer (20 mM Tris-HCl pH 8.1, 2 mM EDTA, 0.5 M NaCl, 0.1% SDS, 1% Triton X-100), followed by lithium chloride buffer (10 mM Tris-HCl pH 8.1, 1 mM EDTA, 0.25 M LiCl, 1% IGEPAL and 1% deoxycholic acid) and TE buffer (10 mM Tris-HCl pH 8.1, 1 mM EDTA).

Chromatin-Antibody complexes from beads were incubated at room temperature for 10 min on magnetic stand and eluted using 100 µl elution buffer (0.1 M NaHCO_3_, 1% SDS). Cross-links were reversed by heating the eluate overnight at 65°C in 5 M NaCl solution, samples were treated with proteinase K (Thermo Fisher Scientific, 26160) for 1 h at 55°C, and DNA was purified using QIAQuick PCR purification kits. Libraries were prepared using ThruPLEX DNA-seq kits (Rubicon Genomics, R400427) and sequenced on an Illumina HiSeqX instrument.

### **Generation of Neurog3 inducible hISC organoids with a fluorescent *ASCL1* reporter**

Homology-directed repair (HDR) donor sequences for in-frame tagging of ASCL1-mNeonGreen and ASCL1-mTurquoise2 were constructed using Gibson Assembly. DNA segments containing 300-bp homology arms 5’ and 3’ to the *ASCL1* stop codon were obtained as gBlocks (Integrated DNA Technologies) with a 21-bp linker sequence (see Table S6). Fluorescence reporter cDNAs, mNeonGreen and mTurquoise2, were amplified by PCR (Q5 Hot Start High-Fidelity DNA Polymerase, M0493S) from Addgene plasmid #98886, with the 21-bp linker sequence at each end. The DNA segments and PCR-amplified reporters were ligated using standard protocols (Gibson Assembly Cloning kit, NEB, E5510S) to create HDR donor products with the sequence 5’ *ASCL1* homology arm + linker + mNeonGreen/mTurquoise2 + linker + 3’ *ASCL1* homology arm (Table S6). These HDR donor products were used as templates for high-throughput PCR amplification (Q5 Hot Start High-Fidelity DNA Polymerase, M0493S). The amplified HDR donor template was confirmed by gel electrophoresis and Sanger sequencing.

To generate *ASCL1* reporter organoid lines, we introduced Cas9:sgRNA ribonucleoprotein complex (RNP) and the HDR template together into hISC^NEUROG3^ by nucleofection. The RNP complex was prepared by incubating sgRNA CCGAGCCCCTCAGAACCAGT, which targets the *ASCL1* stop codon, with spCas9 2NLS (Synthego) at 1:9 ratio in nucleofection solution (Amaxa P3 primary Cell 4D-Nucleofector X kit, V4XP-3012) at room temperature for 10 min, then the HDR donor template was added for 2 min. hISCs were dissociated with TrypLE, centrifuged at 300 *g* for 5 min, washed once with PBS, resuspended in the RNP complex/HDR donor template mixture, and nucleofected (4D-Nucleofector X Unit, Lonza, AAF1003X) using program FF137. Cells were then transferred to pre-warmed hISC cell culture medium, seeded on dishes coated with inactivated MEFs, incubated at 37°C in a 7.5% CO_2_ incubator for 3-4 days, dissociated using TrypLE, and split 1:2 as described above. One aliquot was seeded for clonal expansion and the other was used to extract genomic DNA using Quick-DNA^TM^ Microprep kit (Zymo Research, D3020). Primers ASCL1F and mNeonGreenR or ASCL1F and mTurquoiseR (Table S6) were used to amplify the targeted region (Phire^TM^ Tissue Direct PCR Master Mix, F-170S), purified (Qiagen PCR purification kit, 28006), and sequenced to confirm in-frame integration at the 3’ terminus of *ASCL1*.

### Live imaging of ASCL1-reporter hISC^NEUROG^^3^ cells

hISC^NEUROG3^ cells carrying a fluorescent *ASCL1* reporter were seeded on 1.5 mm glass-bottom 6-well dishes (Cellvis, P06-1.5H-N) coated with inactive MEFs and incubated at 37°C in a 7.5% CO_2_ incubator. To reduce background fluorescence, phenol-free cell culture medium was applied before imaging. Cells were imaged starting 72 h after NEUROG3 induction and images were acquired on THUNDER Imager 3D Cell Culture microscope (Leica Microsystems) using a 20x (0.8 NA) air objective. mTurquoise2, mNeonGreen and mCherry fluorescence images were acquired using detectors for DAPI, fluorescein, and rhodamine, respectively. LED intensity and exposure times for each channel were systematically minimized. In each experiment, 20 regions of interest were selected based on nuclear expression of mNeonGreen or mTurquoise2 and images were captured for 12 h at 10-min intervals between recorded z-stacks of 35 µm thickness. To track cells over time, their boundaries were acquired using the brightfield channel.

### CUT&RUN

CUT&RUN was performed as described previously.^105^ Briefly, cells were dissociated 96 h after NEUROG3 induction, and sorted by FACS to collect 5×10^5^ live (DAPI^-^) mCherry^+^ cells, hence excluding MEF contaminants. Cells were pelleted at 600 *g* for 5 min at 4°C and resuspended in ice-cold nuclear extraction buffer (20 mM HEPES, 100 mM KCl, 0.5 mM spermidine, 0.1% triton X-100, 20% glycerol and 1x Roche Complete Protease Inhibitor EDTA-free), followed by binding of isolated nuclei to Concanavalin A-coated magnetic beads (Polysciences, 86057-3). To chelate divalent cations, bead-bound nuclear slurry was incubated for 5 min with blocking buffer (20 mM HEPES, 150 mM NaCl, 0.5 mM spermidine, 0.09% BSA and 1x Roche Complete Protease Inhibitor EDTA-free) containing 2 mM EDTA. The slurry was then incubated overnight with ASCL1 (Abcam, ab74065, 1:100), NEUROD1 (Cell Signaling Technology, 4373, 1:100) Ab or normal rabbit IgG (negative control, Millipore, 12-370) diluted in blocking buffer containing 0.05% digitonin (Promega, G9441) on a rotating platform at 4°C. The slurry was then washed and coupled with Protein A-conjugated micrococcal nuclease (pA-MNase; 1:200 dilution of a 140 µg/ml stock kindly gifted by S. Henikoff, University of Washington) for 60 min at 4°C. The slurry was washed again to remove unbound MNase and equilibrated in an ice-water bath for 5 min before addition of 2 mM CaCl_2_ to activate MNase and digested nuclease bound regions for 60 min in the ice-water bath. The enzyme was then inactivated in stop buffer (200 mM NaCl, 20 mM EDTA, 4 mM EGTA, 50 µg/ml RNase A and 25 µg/ml glycogen) and cleaved DNA fragments were released by incubation for 30 min at 37°C, extracted with phenol-chloroform, and precipitated with ethanol. Extracted fragments were used to prepare libraries (NEBNext Ultra II DNA Library Prep Kit, NEB, E7645), which were sequenced on a HiSeq-X instrument to generate 10-15 million paired-end 150-bp reads (Novogene). SEACR was used to call peaks^96^ and the computational analysis is described below.

### Targeted gene editing of hISC^NEUROG^^3^ cells

We edited the *HES6*, *ASCL1*, and *NEUROD1* loci using SpCas9 and sgRNAs (Synthego) that were designed using CRISPick (https://portals.broadinstitute.org/gppx/crispick/public). RNP complex was prepared by incubating spCas9 2NLS (Synthego) with sgRNA for the targeting locus (Table S6) at 1:9 ratio in the nucleofector solution (Lonza P3 Primary Cell Solution Box, PBP3-00675) for 10 min at room temperature. Cells were dissociated with TrypLE, centrifuged at 300 *g* for 5 min, washed with PBS, resuspended in the RNP complex mixture (Amaxa^TM^ P3 primary Cell 4D-Nucleofector X kit, V4XP-3012), and nucleofected with Lonza 4D-Nucleofector X Unit (Lonza, AAF1003X) using program FF137. Cells were then transferred to pre-warmed hISC cell culture medium and seeded on dishes coated with inactive MEFs, incubated at 37°C in 7.5% CO_2_ for 3-4 days, dissociated using TrypLE, and split 1:2 as described above. One aliquot was seeded for clonal expansion and the other was used to extract genomic DNA using Quick-DNA^TM^ Microprep kit (Zymo Research, Cat. No. D3020). The targeted region was PCR amplified (Phire^TM^ Tissue Direct PCR Master Mix, F-170S) and the amplicons were sequenced (Azenta/GENEWIZ, South Plainfield, NJ) to assess targeting efficiency.

## QUANTIFICATION AND COMPUTATIONAL ANALYSIS

### Bulk and single-cell (sc) RNA sequencing

Bulk RNA-seq data were aligned to human genome version hg19 and processed using Viper with default parameters to generate raw counts.^101^ Raw counts were normalized and differential gene expression was determined using DESeq2 (Ref.^88^) in R software (R Core Team, version 3.6.3).^95^ Plots were generated using the ggplot2 package in R.^89^

Fastq files from scRNA-seq were aligned to hg19 using CellRanger 3.0.2 (10x Genomics) with default parameters. Quality control, normalization, and clustering were performed in R version 3.6.3 (Ref.^95^) using Seurat package v3.2.3 (Ref.^98^). CellRanger output objects from different days after NeUROG3 induction were merged to create Seurat objects, excluding cells with UMI counts <4,000, expression of <2,000 genes, low RNA complexity (log_10_ genes per UMI <0.8), and mitochondrial gene fraction >0.5. ‘CellCycleScoring’ function was used to infer cell cycle status for each cell. ‘SCTransform’ was used to normalize the merged Seurat object; unwanted sources of variation, such as mitochondrial gene expression and differences between S and G2/M phase scores, were regressed out. The merged object was integrated using ‘FindIntegrationAnchors’ and ‘IntegrateData’ functions in Seurat. Principal component analysis (PCA) dimensionality reduction was performed on the scaled data by ‘RunPCA’. Uniform Manifold Approximation and Projection (UMAP) was performed using 1:50 PCA dimensions to place cells in similar local neighborhood together in two-dimensional space. The same number of PCs was used to construct a K-nearest neighbor (KNN) graph using the ‘FindNeighbors’ function. We clustered cells using the ‘FindClusters’ function at 2.4 resolution, which implements an optimized Louvain algorithm to iteratively group cells together by nearest-neighbor modules. To identify markers enriched in each cluster, we used the ‘FindAllMarkers’ function with default parameters and merged substantively similar clusters. The stem cell cluster (ISCs) in the UMAP was defined based on expression of stem cell markers *LGR5*, *MYC* and *FOXQ1*, which were highly enriched in bulk RNA-seq data from untreated (day 0) hISC. Hormone-expressing cells were annotated based on expression of classic hormone genes. Other clusters were named according to the highest and most selectively expressed TF gene (Table S4).

### Bulk and single-cell (sc) ATAC-sequencing

Bulk ATAC-seq data were aligned to human genome version hg19 using Bowtie2^84^ and PCR duplicates were filtered out. Peaks were identified using MACS2^94,106^ with the q-value cut off 0.001. ATAC peaks from each time point were concatenated and merged for overlapping regions using BEDTools^83^ to give a set of ∼190,000 accessible regions that were used to calculate the Euclidean distance. Promoters (±1 kb from TSS) were discarded to define enhancers and a pairwise differential analysis of each time point identified 82,002 differential enhancers during the time course were grouped into 5 clusters (ISC, early, mid, late, and terminal). Raw signals from individual files were used to create Bigwig files and were normalized using haystack v0.5.5 (Ref.^91^) followed by deeptools2 v3.4.3 (Ref.^87^) to plots heatmaps in descending order of ATAC sites ± 3.0 kb from center of ATAC across each cluster and to visualize read distribution on Integrated Genomics Viewer v2.5.0 (Robinson et al 2011). The same signals were used to calculate the aggregate profile plots for regions overlapping ASCL1 and NEUROD1 binding. Motifs enriched were identified using the HOMER de novo algorithm v4.11 (Ref. ^90^).

Sequenced scATAC-seq libraries from each time point were aligned to human genome version hg19. Transposase cut sites and peak accessibility were identified using the CellRanger ATAC pipeline v1.1.0. The output data from different days was merged and used to perform quality control, normalization, and clustering using the Signac package v1.7.0 (Ref.^107^) in R version 3.6.3. Cells with >2,000 peak fragments, fraction of reads in peak (FRiP) >0.4, genome blacklist ratio <0.005, mononucleosome to nucleosome-free ratio <4 and transcription start site enrichment (TSS Enrichment, defined by ENCODE as a signal-to-noise metric) score >2 were retained. Signac was used to normalize the merged object by running term frequency-inverse document frequency (TF-IDF) and singular value decomposition (SVD) on the TF-IDF normalized matrix to reduce dimensionality by latent semantic indexing (LSI). Non-linear reduction was performed with UMAP using 2:30 LSI dimensions and the same number of dimensions was used to construct a KNN graph. Graph based clustering was performed with the Smart Local Moving algorithm using ‘FindClusters’ at 2.4 resolution. To group scATAC-seq clusters based on our previously annotated scRNA-seq clusters, we used ‘FindTransferAnchors’ and ‘TransferData’ functions to predict correspondence between the two datasets. Once clusters were annotated based on scRNA-seq properties, peaks were identified using the ‘CallPeaks’ function in MACS2 (Ref.^106^) with default parameters and the following additional arguments: −m 10 30 −q =0.001, effective.genome.size =2.7e+09. Peaks overlapping with annotated blacklist regions in hg19 were removed. The resulting peak set for each cell was quantified using the ‘FeatureMatrix’ function and ATAC features enriched in each cluster were identified using the ‘FindAllMarkers’ function, both in Signac. mRNA expression and chromatin access in corresponding clusters were correlated using the GeneOverlap package and TF motif scores were calculated using chromVAR package implemented in Signac.

### Pseudotime trajectories

Gene expression over pseudotime was computed for standalone scRNA-seq data using the Monocle3 package^38^ in R version 3.6.3 (Ref.^95^). Monocle3 constructs initial trajectory inference using the UMAP dimensionality reduction and refines it with a learning principal graph to compute pseudotime values and we used the ‘learn_graph’ function to infer trajectories based on clusters derived from the Seurat pipeline. The ‘use_partition’ parameter was set to TRUE to ensure that the trajectory would partition into branches and the ‘minimal_branch_len’ and ‘geodesmic_ratio’ were set 4 and 0.4, respectively, to control branch lengths and trajectory connections. The ‘order_cells’ function was used for pseudotime ordering, with the ISC cluster defined as the root. Cells were ordered along the trajectory graph based on their pseudotime, representing the progression along the imputed cellular trajectory.

For multiome-sequencing, pseudotime was calculated using MIRA.^42^ Both gene expression and ATAC data from multiome-sequencing were used as inputs. MIRA uses the Palantir algorithm to assign each cell a pseudotime value based on the undirected joint KNN graph showing cells in similar multimodal states. For any selected cellular state, Palantir transforms the undirected joint KNN graph into a directed transport map that travel backwards relative to pseudotime values to track the ancestral origin of the cellular state chosen and assigns each cell a probability of reaching the terminal state. MIRA then uses Palantir terminal fate probabilities to calculate the cellular trajectories and branch points.

### ChIP-seq analyses

ChIP-seq reads were aligned to the hg19 using Bowtie2 version 2.4.1 (Ref. ^84^) to generate bam files and peaks (q < 0.001) were called using MACS2 (Ref.^94^). Bam files were converted into signal files (bigWig) and quantile normalized with haystack v0.5.5 (Ref. ^91^) using 50-bp window across the genome followed by deepTools2 v3.4.3 (Ref.^87^) and were used to calculate the aggregate profile plots and generate heatmaps.

### Single-cell Multiome analysis for gene expression and open chromatin

Sequenced scMultiome-seq libraries from each time point were aligned to human genome hg19 and transposase cut sites and peak accessibility were identified using CellRanger ARC pipeline v1.0.0. The output data were used to perform quality control, normalization, and clustering using Seurat package v4.0.6 (Ref.^97^) and Signac package v1.7.0 (Ref.^107^) in R version 4.1.1.

CellRanger output objects from different days were merged to create a Seurat object, excluding low-quality cells with UMI counts <1,000, expression of <1,000 genes, low RNA complexity (log_10_ genes per UMI <0.8), peak fragments <2,000, FRiP score <0.4, mononucleosome to nucleosome-free ratio >4, and TSSEnrichment score <2. RNA and ATAC data were normalized as described for standalone scRNA and scATAC. To represent both gene expression and ATAC measurements, we computed a joint neighbor graph using the weighted nearest neighbor approach, implemented using ‘FindMultiModalNeighbors’ and ‘RunUMAP’ functions in Seurat v4. To annotate cell types, we transferred cell labels from standalone scRNA data using ‘FindTransferAnchors’ and ‘TransferData’ functions in Seurat. Cell cluster identities were corroborated unsupervised clustering based on MIRA topics.^42^ Once clusters were thus annotated, peaks were identified using the ‘CallPeaks’ function in MACS2 (Ref.^106^), as described above for standalone scATAC-seq. To plot correlation between peak accessibility, motif enrichment and gene expression we used ArchR package.^82^

For CRISPR-edited knockout experiment, CellRanger output objects from scrambled gRNA (day 3 n=2, day 6 n=2), *ASCL1*-null (days 3 and 6), *HES6*-null (days 3 and 6), *NEUROD1^DBD^*, and *NEUROD1^TAD^* (both only day 6) were used as input. For *HES6*-null and *NEUROD1^DBD^*, we only retained cells with UMI counts >1,000, expression of >500 genes, high RNA complexity (log_10_ genes per UMI) >0.8, peak fragments >2,000, FRiP score >0.2, mononucleosome to nucleosome-free ratio <4, and TSS enrichment score >2. Scrambled gRNA and TF gene-specific knockout cells were normalized as described for standalone scRNA and scATAC and integrated using ‘SelectIntegrationFeatures’, ‘FindIntegrationAnchors’ and ‘IntegrateData’ with default parameters in Seurat v4. Integrated datasets were scaled and cells with differences between S and G2/M phase scores were regressed out. Dimensions were reduced by PCA and UMAP to place cells together in two-dimensional space and cells were annotated according to standalone scRNA classifications.

### Quantitation of ASCL1 live cell imaging

Individual cells were tracked, and fluorescence signals quantified with Imaris software (Bitplane, RRID:SCR_007370) and ImageJ (RRID:SCR_002285) using the ‘Circadian Gene Expression’ plugin (http://bigwww.epfl.ch/sage/soft/circadian/). mCherry and brightfield images were used to detect and track cell motion. We obtained maximum intensities for each z-plane from >20 cells for mNeonGreen and mTurquoise2 fluorescence levels, quantified at 10-min intervals over 12 h periods of imaging. To exclude noise, fluorescence counts were smoothened to identify peaks and troughs during this 12 h period. One oscillation cycle for fluorescent ASCL1 was defined as the interval from one peak or trough to the next.

### CUT&RUN analyses

Sequenced reads for ASCL1 and NEUROD1 were processed using CUT&RUNTools^76^ to trim, align and filter fragments to identify those likely to contain TF binding sites, i.e., <120 bp in length. Briefly, paired-end 150-bp reads were trimmed using Trimmomatic and Kseq and aligned to human genome hg19 using Bowtie2 version 2.4.1. Peaks were called using SEACR (Sparse Enrichment Analysis for CUT&RUN)^96^ using default parameters; potentially artifactual binding sets were excluded by intersecting a set of concordant peaks from at least 3 replicates for each TF with a union set of ATAC-seq peaks. These SEACR-called peaks were used as inputs for motif and footprint analyses within CUT&RUNTools. Raw signals from mapped reads for IgG, ASCL1 and NEUROD1 CUT&RUN were used to create bigwig files that were normalized for sequencing depth and heatmaps were plotted over regions grouped as shared or enriched for either ASCL1 or NEUROD1 binding using deeptools2 v3.4.3 (Ref.^87^) in descending order ± 1.5 kb from center of binding. The same signals were used to calculate the aggregate profile plots.

### Quantitation of Crispr/Cas9 knockout efficiency

Amplicon sequenced reads were processed using CRISPResso2 (Ref. ^108^), which quantifies the knockout efficiency based on the fraction of reads showing deletions, substitutions, or insertions within the target region. Most lesions were small deletions.

## QUANTIFICATION AND STATISTICAL ANALYSIS

Statistical analyses were performed using GraphPad Prism 9 or R. Tests and individual biological or technical replicates for each experiment are specified in figure legends. Statistical results are presented as averages, with individual values shown. Error bars represent the SEM. Graphical illustrations were created using BioRender.com.

---

**Figure S1.**
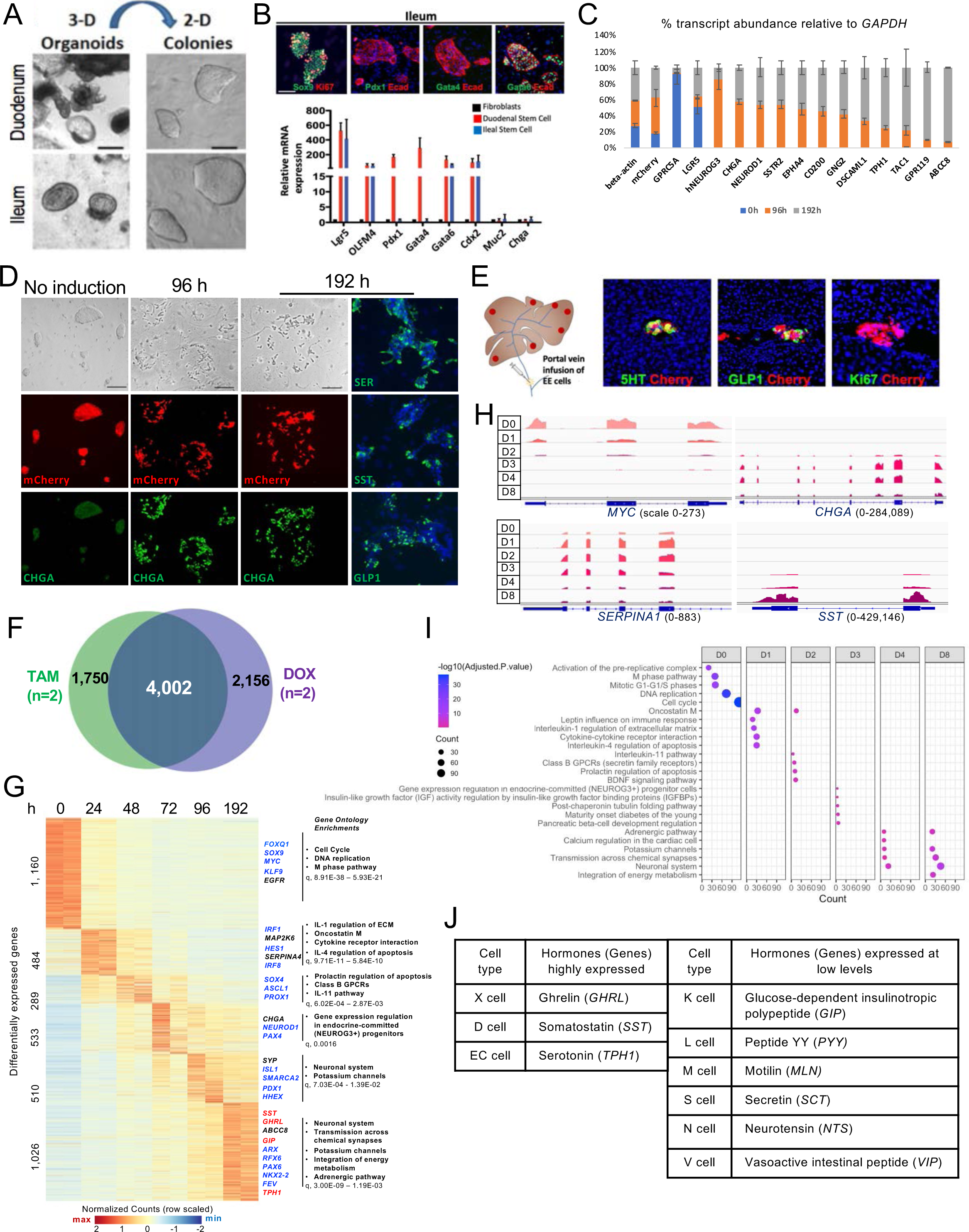
Biology, regional identity, and EEC differentiation of hISC^NEUROG3^ cells. **Related to** Figure 1. (A) Human intestinal biopsies, first cultured as 3D organoids in Matrigel, were subsequently expanded as 2D colonies on tissue culture dishes coated with a feeder layer of mitotically inactive mouse embryonic fibroblasts (MEFs). Scale bars 100 µm. (B) Ileal hISCs express SOX9 (an intestinal stem-cell marker), KI67 (a proliferative cell marker), GATA6 (a distal intestine marker), and E-cadherin (Ecad, an epithelial marker) but they lack proximal intestinal markers PDX1 and GATA4. Gene expression, determined by qRT-PCR and displayed relative to human fibroblasts, shows enrichment of broad intestinal markers *GATA6* and *CDX2* and of stem cell markers *LGR5* and *OLFM4* in both duodenal and ileal cells, and of proximal intestinal markers *PDX1* and *GATA4* only in duodenal cells. *CHGA*, a marker of differentiated EECs, was absent from uninduced hISCs. (C) Transcript abundance (qRT-PCR, shown relative to *GAPDH*) reveals uniform expression of ý-actin and mCherry across time. Stem-cell markers *GPRC5A* and *LGR5* were abundant in untreated cells (0 h) and EEC markers (native *NEUROG3, CHGA, NEUROD1, SSTR2, EPHA4, CD200* and *GNG2*) were present by 96 h. Late EEC markers (*DSCAML1, TPH1, TAC1, GPR119* and *ABCC8*) were enriched at 192 h. (D) Bright-field and fluorescence microscopy of uninduced hISCs reveal mCherry expression in compact colonies that lack CHGA (scale bars 100 µm). Following 48 h of Tam exposure, mCherry expressing hISC^NEUROG^^3^ cells were dispersed and expressed CHGA by 96 h. Induced cells expressed serotonin (SER, 5HT, marks EC cells), SST (somatostatin, D cells) and GLP-1 (glucagon-like peptide 1, L cells). (E) Tam-induced hISC^NEUROG^^3^ cells were infused into the portal veins of immunodeficient NSG mice. Two months later, recipient livers contained numerous clusters of non-proliferating mCherry^+^ cells that expressed 5HT and GLP-1. (F) Multi-pairwise analysis of bulk RNA-seq samples at 0, 24, 48, 72, 96 and 192 h after initial Tam or doxycycline (Dox) exposure revealed 4,002 genes differentially expressed between any two time points in two independent ileal hISC^NEUROG^^3^ lines (n=2 replicates each), one dependent on tamoxifen and other dependent on doxycycline to activate NEUROG3. (G) Dynamic expression of 4,002 genes over the course of EEC differentiation and maturation. Intestinal stem cell (ISC) markers, e.g., *FOXQ1, SOX9, MYC* and *KLF9*, decline soon after NEUROG3 activation, giving way to EEC markers such as *SOX4, PROX1, NEUROD1, CHGA and SYP* between 72 h and 96 h, followed by enrichment of hormone genes such as *SST, GHRL, GIP* and *TPH1* at 192 h. Selected Gene Ontology (GO) terms enriched among genes differentially enriched on different days are listed to the right. (H) Integrative Genomics Viewer (IGV) tracks showing bulk RNA-seq reads from representative ISC (*MYC*), intermediate cell (*SERPINA1*), and mature EEC (*CHGA* and *SST*) genes. (I) Over the course of EEC differentiation, GO terms representing differentially expressed genes (Figure S1G) show rapid loss of proliferative cell properties, early interim immune signaling, and subsequent acquisition of neurosecretory features. (J) Mature EEC types and their known specific hormones.

**Figure S2.**
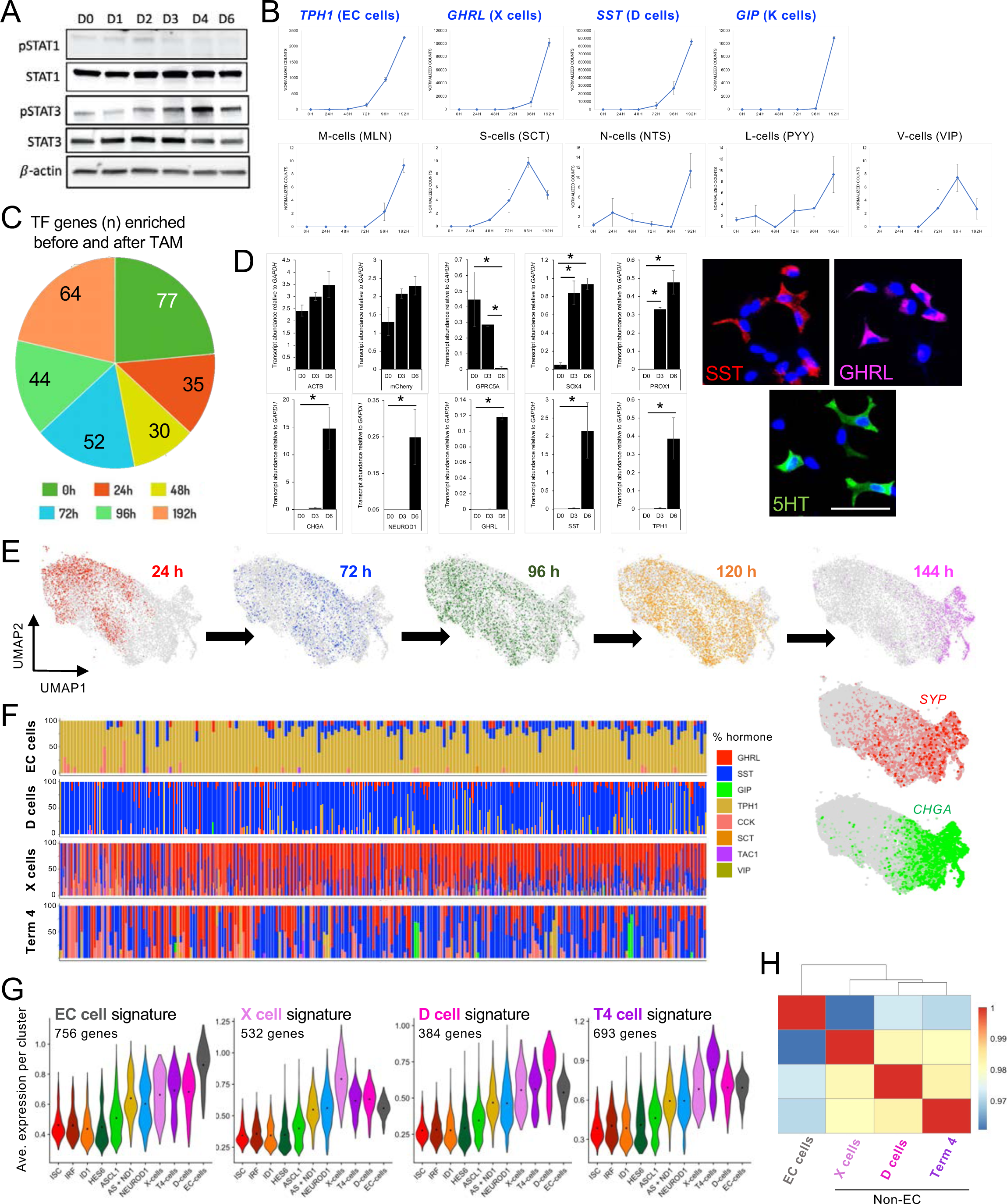
Characterization of human EECs differentiated *in vitro*. Related to Figure 1. (A) Immunoblots of protein lysates 0, 24, 48, 72, 96 and 144 h after hISC^NEUROG^^3^ cell treatment with Tam reveal transient pSTAT1 and pSTAT3 activation. Total STAT1 and STAT3 levels were unaltered; ß-actin serves as a loading control. (B) During EEC differentiation, transcripts for hormones corresponding to specific EEC types (normalized bulk RNA-seq counts, DEseq2) increased late. The 4 most abundant products correspond to EC, X, D, and K cells; others were expressed at substantially lower levels. (C) Numbers of transcription factor (TF) genes enriched on successive days of Tam-induced EEC differentiation of hISC^NEUROG^^3^ cells. (D) Left: qRT-PCR analysis (values normalized to *GAPDH*) 72 h and 144 h after NEUROG3 activation shows substantial enrichment of terminal markers (*CHGA*, *NEUROD1*, *GHRL*, *PAX6* and *SST*) at 144 h, compared to early (GPRC5A) and intermediate (*SOX4*, *PROX1*) markers, which decline or appear by 72 h (means ±SEM, n=3 independent experiments, * p <0.05 from unpaired t-test). Right: Immunofluorescence for SST, GHRL, and 5HT at 144 h (scale bar 50 µm). Thus, 144 h is sufficient time to investigate lineage determinants. (E) Integrated UMAP from the full course of EEC differentiation showing the positions of cells isolated 24 h, 72 h, 96 h, 120 h and 144 h after initial Tam exposure. Differentiation is not strictly synchronized. At 96 h or 120 h, cells represent a spectrum of pre-terminal states, but by 144 h, nearly all remaining cells are readily recognized as terminal X, D, T4 or EC cells. Feature plots below reveal that *SYP* (red) precedes *CHGA* (green) expression. (F) Relative levels of individual hormone mRNAs in the pool of all hormone transcripts in individual cells of each terminal cluster. Each column in a block represents a single cell. (G) mRNA signatures that distinguish terminal human EC, X, D, and T4 cells differentiated *in vitro* (quantitation of enriched genes listed in Table S3). (H) Pearson correlations among all transcripts detected by scRNA-seq. The largest distinction is between EC and non-EC (X, D, and T4) cell types.

**Figure S3.**
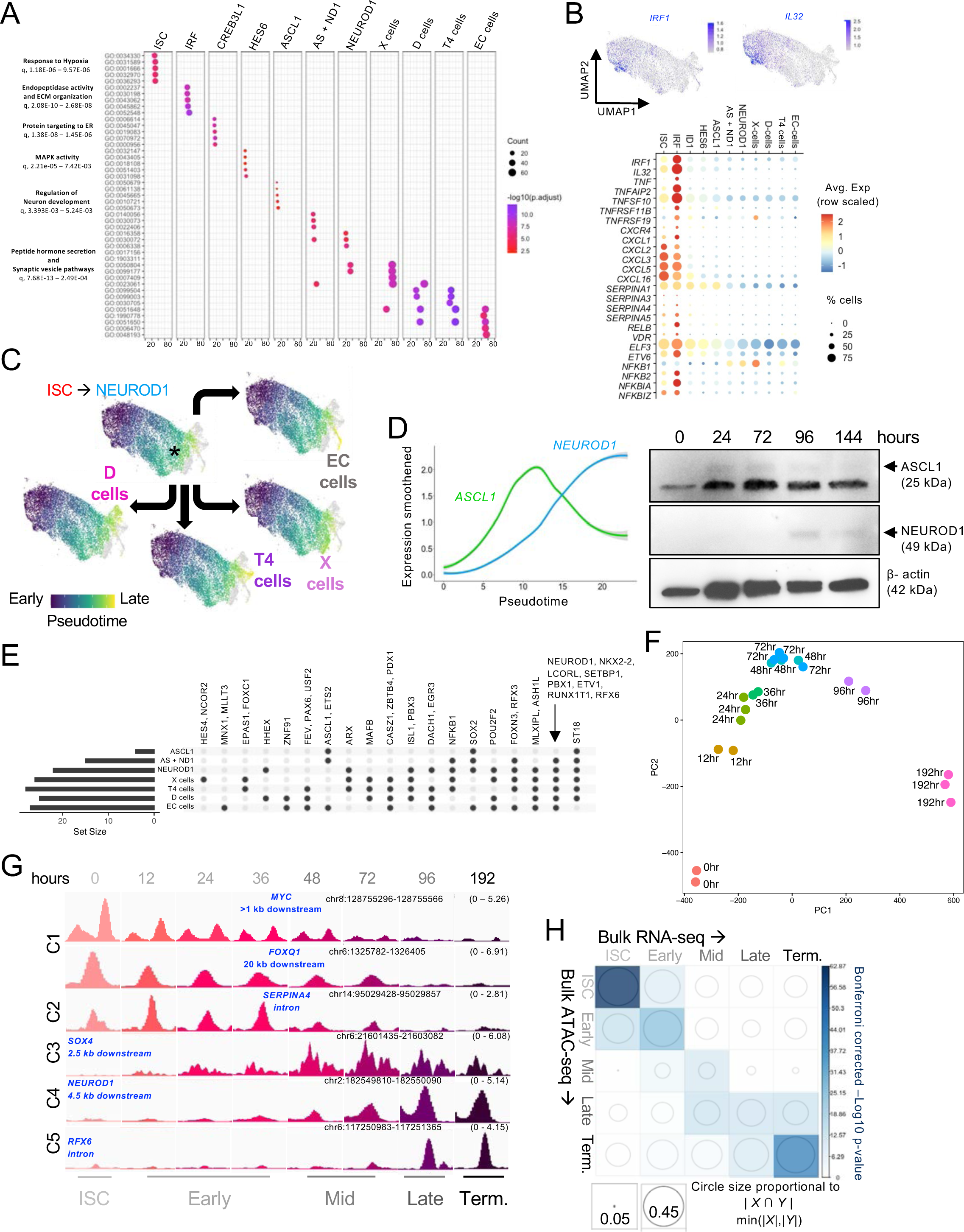
Characterization of EEC types from scRNA-seq data. Related to Figure 2. (A) Gene ontology (GO) term enrichments for genes differentially expressed in 11 cell clusters (Figure 1F) recognized along the human EEC differentiation trajectory. (B) Top: *IRF1* and *IL32* (representative genes) expression projected onto the integrated UMAP of human EEC differentiation. Bottom: Selectively enriched expression of genes related to interferon signaling in the early cell cluster dominated by *IRF1* expression. (C) Pseudotime trajectories derived from Monocle3 analyses of scRNA-seq data. Early stages are represented in dark, and later stages in light, colors. After following a common path from ISCs to *NEUROD1^+^* precursors, the trajectory reveals divergence to D, T4, X, and EC cells. (D) Left: Smoothened plots of pseudotemporal *ASCL1* and *NEUROD1* mRNA expression. Right: Immunoblots showing the appearance of ASCL1 before NEUROD1 in bulk protein lysates harvested on different days after initial Tam exposure of hISC^NEUROG^^3^ cells. (E) Unique and combinatorial TF gene expression in each cellular state along the human EEC trajectory (see Figure 2B). (F) Principal component (PC) analysis reveals concordance among replicate ATAC-seq libraries from cells isolated on different days after NEUROG3 (Tam) induction. (G) IGV tracks from bulk ATAC-seq of representative enhancers selectively accessible in ISCs (C1: *MYC* and *FOXQ1*), transiently accessible early in EEC differentiation (C2 and C3: *SERPINA4* and *SOX4*), or only accessible late in EEC maturation (C4 and C5: *NEUROD1* and *RFX6*). (H) Relation between gene expression (bulk RNA-seq, columns) and genes located near sites of differential open chromatin (bulk ATAC-seq, rows) in EEC differentiation stages classified as ISCs (untreated cells, 0 h), early (12, 24, and 36 h), mid (48 and 72 h), late (96 h), or terminal (192 h). Circle diameters represent the fractional overlap between ATAC peaks and expression levels of the nearest gene within 25 kb, computed at the gene level for each stage. The color scale represents Bonferroni-corrected *P* values from a hypergeometric test.

**Figure S4.**
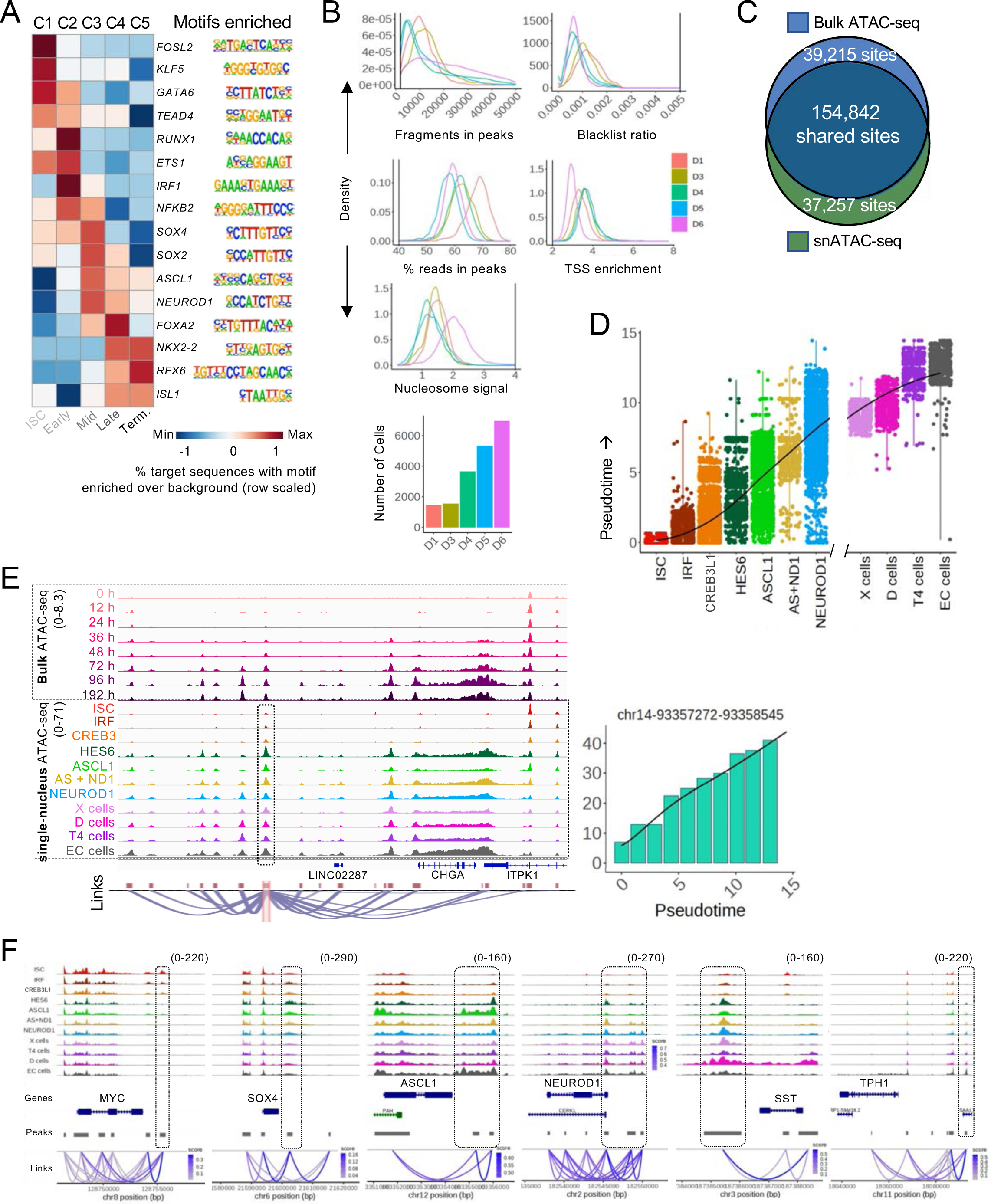
Chromatin dynamics during human EEC differentiation. Related to Figure 2. (A) Enriched DNA sequence motifs identified *de novo* in different ATAC-seq clusters (C1 to C5) during EEC differentiation. Sites selectively accessible in ISCs were enriched for motifs that correspond to ISC-enriched TFs, such as FOSL2 and KLF5. After NEUROG3 induction, early-accessible (12 h to 36 h) sites were enriched for inflammatory TFs ETS1, IRF1 and NFKB2, followed by motifs for EEC factors SOX4, SOX2 and ASCL1 and NEUROD1, and culminated in motifs for terminal EEC factors such as NKX2-2, RFX6, and ISL1. (B) Quality control metrics for snATAC-seq: Fragments present within peaks, blacklist ratios, fractions of reads in peaks (FRIP scores), transcriptional start site (TSS) enrichment, and nucleosome signals. Each curve represents the cells isolated on a specific day after initial Tam exposure. Cell numbers meeting our stringent criteria are shown for each day. (C) Overlap between ATAC-seq peaks called from bulk and single-nucleus (sn) analysis. (D) Pseudotime trajectory derived from Monocle3 analysis of snATAC-seq scores, revealing a common path from ISCs to *NEUROD1^+^*EEC precursors, followed by divergence into 4 terminal cell types. (E) IGV tracks showing the evolution of chromatin accessibility at the *CHGA* locus in bulk (top, different times, scale 0-8.3) and sn (bottom, pseudotemporal cell clusters, scale 0-71) ATAC-seq. Although snATAC-seq data are sparse in any cell, pseduobulking of clustered cells matches the profiles we observed on different days in bulk ATAC-seq, even at sites with relatively weak signals, hence raising confidence in data quality, cluster assignments, and pseudotemporal order. Right, snATAC signals at one peak (dotted outline) are plotted across pseudotime bins. Below, “links” signify ATAC site co-accessibility, as determined by Cicero analysis.^109^ (F) IGV tracks for peaks across snATAC-seq defined pseudobulk cellular states representing early (*MYC*), mid (SOX4 and *ASCL1*) and terminal EEC types (*NEUROD1*, *SST* and *TPH1*). Peaks downstream of *MYC* progressively relinquish access while a *SOX4*-downstream peak is transiently accessible. Peaks surrounding *ASCL1*, *NEUROD1*, *SST*, and *TPH1* become progressively accessible across all maturing EEC types and remain open even in cells where the gene (e.g., *ASCL1* and *TPH1*) is not expressed.

**Figure S5.**
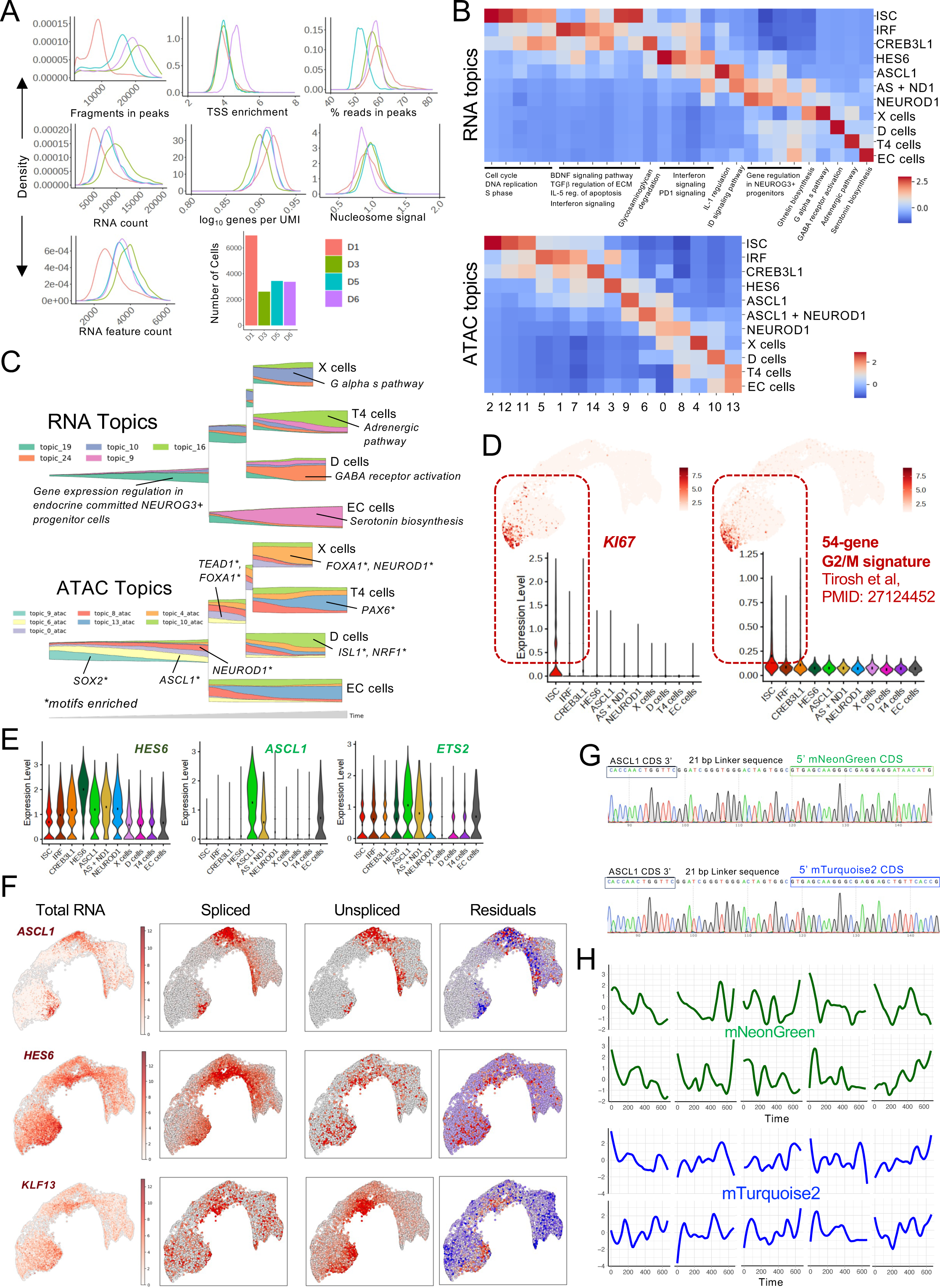
Single-nucleus (sn) Multiome-seq analysis of EEC differentiation and ASCL1 oscillation. Related to Figures 3 and 4. (A) Quality control metrics for snMultiome-seq. For the ATAC modality: fragments in peaks, TSS enrichment, fraction of reads in peaks (FRIP score), and nucleosome signal. For the RNA modality: total RNA counts, RNA feature counts, and RNA complexity (log_10_ genes per UMI). Cell numbers that met stringent criteria and which we considered further are shown for each time between days 1 and 6 after initial Tam exposure on day 0. (B) Using RNA (top) and ATAC (bottom) topics, a probabilistic multimodal model for integrated regulatory analysis (MIRA) clustered cells independent of conventional nearest-neighbor approaches. RNA and ATAC topic-based cell clusters closely matched the clusters defined by scRNA-seq (Figure 1D) and, after transfer of gene anchors, by scATAC-seq (Figure 2D). This correspondence further enhances confidence in the designated interim and terminal EEC states. (C) Stream plots of RNA expression and chromatin accessibility topics across the EEC cell state trajectory. Pathways activated in expression topics and DNA sequence motifs enriched in ATAC topics are indicated. As with stand-alone scRNA-seq and snATAC-seq analyses, the topic-based trajectory indicated a common path from ISCs to *NEUROD1^+^* EEC precursors, followed by divergence into X, T4, D, and EC cells. (D) Projection of *KI67* expression and a 54-gene proliferative gene signature (G2/M score) on the integrated snMultiome UMAP (expression in each of the 11 cell clusters is represented below). Cell replication is confined to ISCs and the earliest EEC precursors and is notably absent from at least the *HES6^hi^* cell state onward, including *ASCL1^+^*precursors. (E) Expression of the *HES6^hi^* cell signature (43 genes) and of *ASCL1* and *ASCL1*-coexpressed gene *ETS2* in each pseudotemporal cell cluster. (F) Feature plots from snMultiome-seq showing the levels of total RNA, spliced and unspliced forms, and unspliced unique transcript count residuals (red = high) for *ASCL1*, *HES6*, and *HES6*-coexpressed gene *KLF13*. Ongoing *HES6* and *KLF13* transcription (unspliced RNA) is evident principally in the *HES6*-dominant cell cluster. (G) DNA sequence confirmation of accurate in-frame engineering of mNG and mT2 fluorescent tags into the 3’ end of the *ASCL1* gene, separated by a 21-bp linker. (H) Quantified time-lapse expression (nuclear fluorescence at 10-min intervals over 12 h starting ∼72 h after Tam exposure) of representative hISC^NEUROG3^ cells with mNG- (top) or mT2- (bottom) tagged ASCL1. Oscillatory periods (inter-peak and inter-trough intervals) in these and additional cells are represented in Figure 4E.

**Figure S6.**
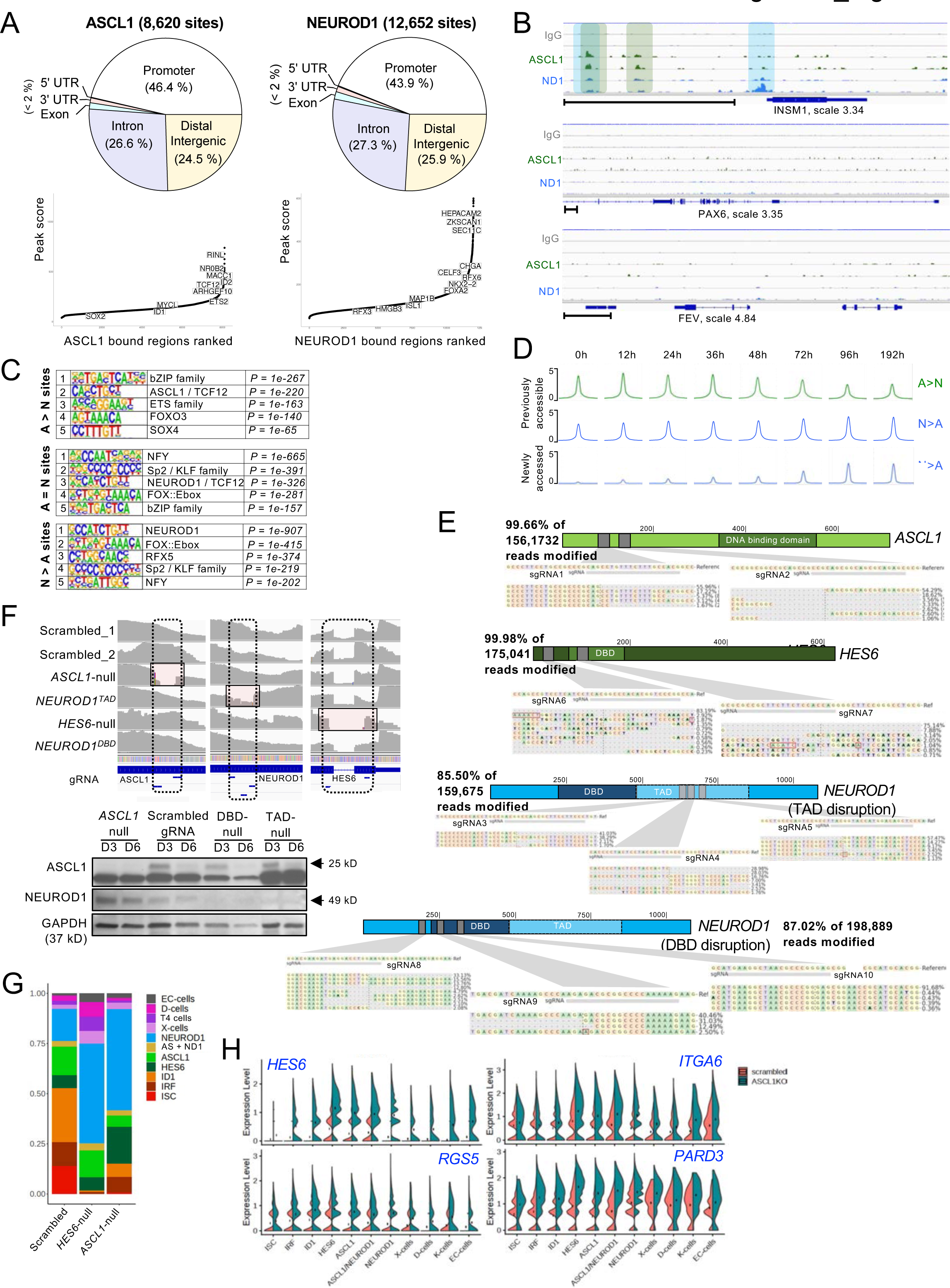
TF binding at EEC *cis*-elements and Crispr/Cas9 editing to generate TF-mutant hISC^NEUROG3^ cells. Related to Figures 5 and 6. (A) Top, ASCL1 and NEUROD1 binding sites identified by CUT&RUN fall mainly in promoters (<2 kb from TSSs) and in presumptive intronic and distal intergenic enhancers. Bottom, Binding sites plotted in rank order of CUT&RUN signal strength highlight the nearest gene highly expressed in *ASCL1^+^*and *NEUROD1^+^* maturing EEC clusters. (B) IGV tracks showing ASCL1 and NEUROD1 binding upstream of *INSM1* and absence of binding surrounding *PAX6* and *FEV* loci, TFs expressed in terminal EECs. (C) DNA sequence motifs most enriched at differentially occupied (A>N and N>A) and shared (A=N) binding sites include E-box variants recognized by ASCL1, TCF12 or NEUROD1. (D) Aggregate ATAC-seq (from bulk data, Figure 2D) signals in previously accessible A>N and N>A binding sites and newly accessed N>A sites over the full course of EEC differentiation. ASCL1-preferred sites gradually relinquish chromatin accessibility, consistent with late drop in *ASCL1* expression, whereas NEUROD1-preferred sites maintain or increase accessibility, in line with persistent *NEUROD1* expression in terminal EEC differentiation. (E) Amplicon sequences across regions targeted by sgRNAs for Crispr/Cas9-mediated disruption of *ASCL1* (null), *HES6* (null) and *NEUROD1* (to generate TAD-null or DBD-null) show >99% and 85-87% efficiency at generating indels (predominantly deletions) around the predicted cleavage sites. (F) Top, aggregated (pseudobulk) snMultiome-seq data showing paucity of RNA reads around the targeted (boxed) *ASCL1, NEUROD1* (TAD or DBD), and *HES6* sites. Bottom, immunoblot confirmation of TF gene disruptions. Protein lysates from control (scrambled gRNA) and CRISPR/Cas9-edited *ASCL1*- and *NEUROD1*-mutant cells 72 h (D3) and 144 h (D 6) after NEUROG3 activation. GAPDH serves as a loading control. (G) Fractions of each cell cluster among all cells derived from control, *HES6*- and *ASCL1*-null cells on the 3^rd^ day after initial Tam exposure. (H) Expression of representative *HES6^hi^* signature genes in all *ASCL1*-null cell derivatives. Data for maturing *NEUROD1^+^* (ND1) cells and their mature descendants, extracted from this panel of all cell clusters, are shown in Figure 6F.

**Figure S7.**
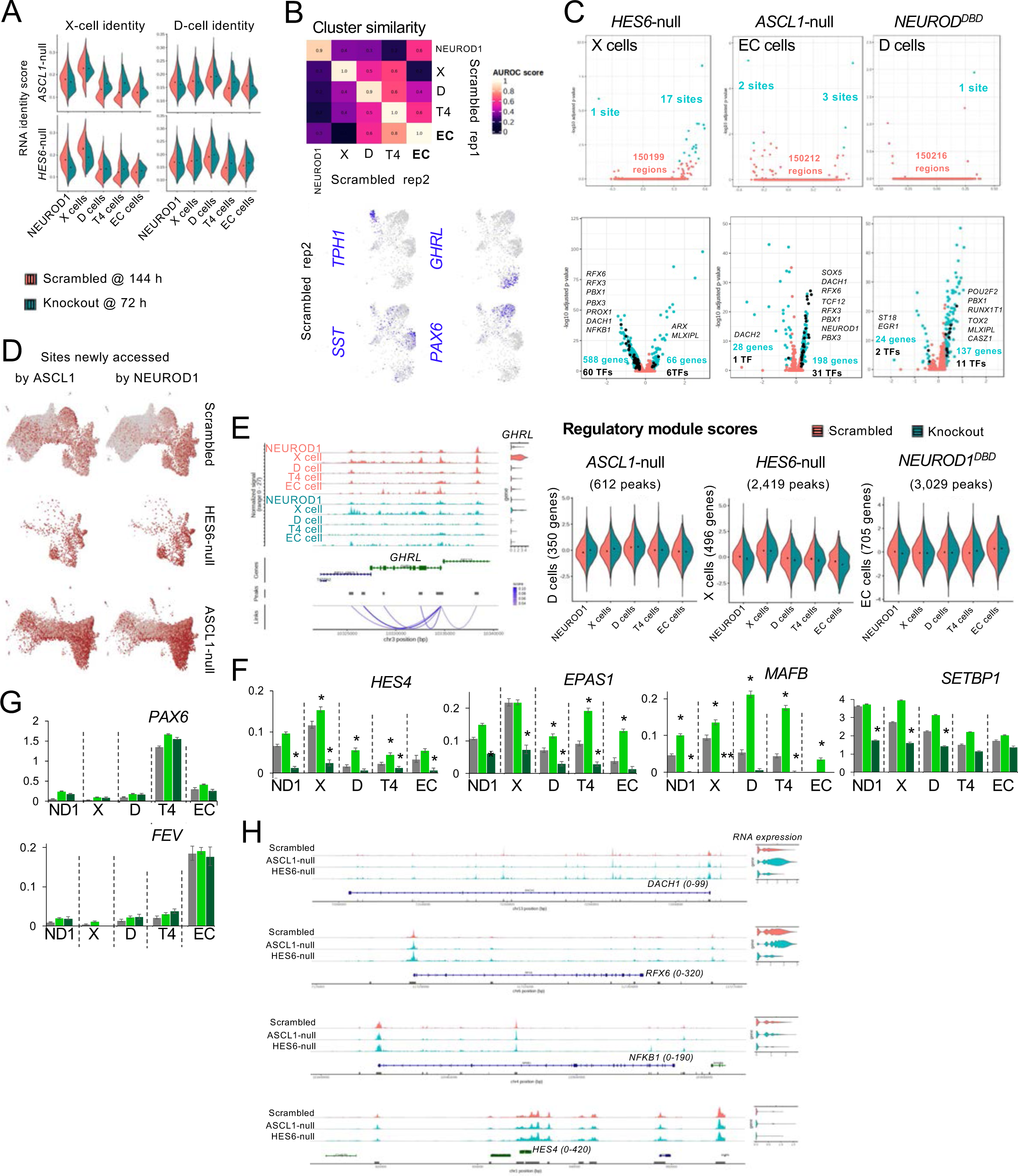
**mRNA and chromatin defects in TF-mutant hISC^NEUROG^**^3^. Related to Figure 7. (A) 72 h after initial Tam exposure, *HES6*-null derivatives show reduced X-cell features and *ASCL1*-null cells hint at enhanced X-cell features in EEC derivatives other than X cells. (B) Cluster similarity between the two scrambled replicates shown using area under the receiver operator characteristic (AUROC) scores (range: 0 to 1, where 1 = perfect similarity, ≥0.9 = high similarity, 0.5 = similarity as good as random, ≤0.3 = high-confidence dissimilarity). *GHRL*, *SST*, *TPH1*, and *PAX6* levels in Scrambled gRNA replicate 2, projected on the UMAP, localize accurately to X, D, EC, and T4 cells, respectively. (C) Differential enhancer accessibility (snATAC, top row of volcano plots) and mRNA expression (snRNA-seq, bottom row) in representative terminal mutant cells, compared in each panel to the corresponding controls. Differences in gene expression arise in the setting of little to no perturbation at >150,000 sites of open chromatin (numbers of statistically significant differences are indicated in teal font). Among differentially expressed genes, those encoding TFs normally expressed in terminal EECs are listed. (D) ASCL1- (left) and NEUROD1- (right) bound enhancers first accessed during EEC maturation (accordingly open only after *ASCL1* or *NEUROD1* expression, Figure 5) are open in *ASCL1*- and *HES6*-null EECs. Thus, both factors are dispensable for their accessibility *per se*. (E) Example of ATAC site-RNA expression associations from which we derived regulatory module scores for CREs in each cell type. D-, X-, and EC scores for chromatin accessibility were unchanged in mutants showing the most perturbed gene expression in D (*ASCL1*-null), X (*HES6*-null), and EC (*NEUROD1^DBD^*) cells. (F) Differential TF gene expression in mutant EECs. Average expression of genes in addition to those shown in Figure 7F. (G) Undisturbed expression of TF genes ordinarily restricted to a single terminal EEC type, e.g., *PAX6* (T4 cells) and *FEV* (EC cells), in *ASCL1*- or *HES6*-null derivatives. (H) IGV tracks showing similar chromatin accessibility in scrambled gRNA-edited (control) and mutant cells at representative TF loci with altered expression in mutant cells. *DACH1* (X-cell marker) and *RFX6* (pan-EEC marker) transcripts were increased in *ASCL1*- and decreased in *HES6*-null derivatives, while X-cell markers *NFKB1* and *HES4* were attenuated in *HES6*-null cells.

## REFERENCES

1. Heintzman, N.D., Stuart, R.K., Hon, G., Fu, Y., Ching, C.W., Hawkins, R.D., Barrera, L.O., Van Calcar, S., Qu, C., Ching, K.A., et al. (2007). Distinct and predictive chromatin signatures of transcriptional promoters and enhancers in the human genome. Nat Genet 39, 311–318. 10.1038/ng1966.

2. Calo, E., and Wysocka, J. (2013). Modification of enhancer chromatin: what, how, and why? Mol Cell 49, 825–837.

3. Wamstad, J.A., Alexander, J.M., Truty, R.M., Shrikumar, A., Li, F., Eilertson, K.E., Ding, H., Wylie, J.N., Pico, A.R., Capra, J.A., et al. (2012). Dynamic and coordinated epigenetic regulation of developmental transitions in the cardiac lineage. Cell 151, 206–220. 10.1016/j.cell.2012.07.035.

4. Li, L., Wang, Y., Torkelson, J.L., Shankar, G., Pattison, J.M., Zhen, H.H., Fang, F., Duren, Z., Xin, J., Gaddam, S., et al. (2019). TFAP2C- and p63-dependent networks sequentially rearrange chromatin landscapes to drive human epidermal lineage commitment. Cell Stem Cell 24, 271–284 e278. 10.1016/j.stem.2018.12.012.

5. Lengerke, C., Grauer, M., Niebuhr, N.I., Riedt, T., Kanz, L., Park, I.H., and Daley, G.Q. (2009). Hematopoietic development from human induced pluripotent stem cells. Ann N Y Acad Sci 1176, 219–227. 10.1111/j.1749-6632.2009.04606.x.

6. Spence, J.R., Mayhew, C.N., Rankin, S.A., Kuhar, M.F., Vallance, J.E., Tolle, K., Hoskins, E.E., Kalinichenko, V.V., Wells, S.I., Zorn, A.M., et al. (2011). Directed differentiation of human pluripotent stem cells into intestinal tissue in vitro. Nature 470, 105–109. 10.1038/nature09691.

7. Yang, N., Ng, Y.H., Pang, Z.P., Sudhof, T.C., and Wernig, M. (2011). Induced neuronal cells: how to make and define a neuron. Cell Stem Cell 9, 517–525. 10.1016/j.stem.2011.11.015.

8. Gribble, F.M., and Reimann, F. (2019). Function and mechanisms of enteroendocrine cells and gut hormones in metabolism. Nat Rev Endocrinol 15, 226–237. 10.1038/s41574-019-0168-8.

9. Martin, A.M., Sun, E.W., Rogers, G.B., and Keating, D.J. (2019). The influence of the gut microbiome on host metabolism through the regulation of gut hormone release. Front Physiol 10, 428. 10.3389/fphys.2019.00428.

10. Wang, J., Cortina, G., Wu, S.V., Tran, R., Cho, J.H., Tsai, M.J., Bailey, T.J., Jamrich, M., Ament, M.E., Treem, W.R., et al. (2006). Mutant neurogenin-3 in congenital malabsorptive diarrhea. N Engl J Med 355, 270–280. 10.1056/NEJMoa054288.

11. Mellitzer, G., Beucher, A., Lobstein, V., Michel, P., Robine, S., Kedinger, M., and Gradwohl, G. (2010). Loss of enteroendocrine cells in mice alters lipid absorption and glucose homeostasis and impairs postnatal survival. J Clin Invest 120, 1708–1721. 10.1172/JCI40794.

12. Talchai, C., Xuan, S., Kitamura, T., DePinho, R.A., and Accili, D. (2012). Generation of functional insulin-producing cells in the gut by Foxo1 ablation. Nat Genet 44, 406–412. 10.1038/ng.2215.

13. Ariyachet, C., Tovaglieri, A., Xiang, G., Lu, J., Shah, M.S., Richmond, C.A., Verbeke, C., Melton, D.A., Stanger, B.Z., Mooney, D., et al. (2016). Reprogrammed stomach tissue as a renewable source of functional beta cells for blood glucose regulation. Cell Stem Cell 18, 410–421. 10.1016/j.stem.2016.01.003.

14. Bany Bakar, R., Reimann, F., and Gribble, F.M. (2023). The intestine as an endocrine organ and the role of gut hormones in metabolic regulation. Nat Rev Gastroenterol Hepatol ePub ahead of print. 10.1038/s41575-023-00830-y.

15. Roth, K.A., and Gordon, J.I. (1990). Spatial differentiation of the intestinal epithelium: analysis of enteroendocrine cells containing immunoreactive serotonin, secretin, and substance P in normal and transgenic mice. Proc Natl Acad Sci USA 87, 6408–6412. 10.1073/pnas.87.16.6408.

16. Beumer, J., Artegiani, B., Post, Y., Reimann, F., Gribble, F., Nguyen, T.N., Zeng, H., Van den Born, M., Van Es, J.H., and Clevers, H. (2018). Enteroendocrine cells switch hormone expression along the crypt-to-villus BMP signalling gradient. Nat Cell Biol 20, 909–916. 10.1038/s41556-018-0143-y.

17. Jenny, M., Uhl, C., Roche, C., Duluc, I., Guillermin, V., Guillemot, F., Jensen, J., Kedinger, M., and Gradwohl, G. (2002). Neurogenin3 is differentially required for endocrine cell fate specification in the intestinal and gastric epithelium. EMBO J 21, 6338–6347. 10.1093/emboj/cdf649.

18. Bjerknes, M., and Cheng, H. (2006). Neurogenin 3 and the enteroendocrine cell lineage in the adult mouse small intestinal epithelium. Dev Biol 300, 722–735. 10.1016/j.ydbio.2006.07.040.

19. Sinagoga, K.L., McCauley, H.A., Munera, J.O., Reynolds, N.A., Enriquez, J.R., Watson, C., Yang, H.C., Helmrath, M.A., and Wells, J.M. (2018). Deriving functional human enteroendocrine cells from pluripotent stem cells. Development 145, dev165795. 10.1242/dev.165795.

20. Naya, F.J., Huang, H.P., Qiu, Y., Mutoh, H., DeMayo, F.J., Leiter, A.B., and Tsai, M.J. (1997). Diabetes, defective pancreatic morphogenesis, and abnormal enteroendocrine differentiation in BETA2/neuroD-deficient mice. Genes Dev 11, 2323–2334. 10.1101/gad.11.18.2323.

21. Larsson, L.I., St-Onge, L., Hougaard, D.M., Sosa-Pineda, B., and Gruss, P. (1998). Pax 4 and 6 regulate gastrointestinal endocrine cell development. Mech Dev 79, 153–159. 10.1016/s0925-4773(98)00182-8.

22. Gierl, M.S., Karoulias, N., Wende, H., Strehle, M., and Birchmeier, C. (2006). The zinc-finger factor Insm1 (IA-1) is essential for the development of pancreatic beta cells and intestinal endocrine cells. Genes Dev 20, 2465–2478. 10.1101/gad.381806.

23. Desai, S., Loomis, Z., Pugh-Bernard, A., Schrunk, J., Doyle, M.J., Minic, A., McCoy, E., and Sussel, L. (2008). Nkx2.2 regulates cell fate choice in the enteroendocrine cell lineages of the intestine. Dev Biol 313, 58–66. 10.1016/j.ydbio.2007.09.047.

24. Beucher, A., Gjernes, E., Collin, C., Courtney, M., Meunier, A., Collombat, P., and Gradwohl, G. (2012). The homeodomain-containing transcription factors Arx and Pax4 control enteroendocrine subtype specification in mice. PLoS One 7, e36449. 10.1371/journal.pone.0036449.

25. Roberts, G.P., Larraufie, P., Richards, P., Kay, R.G., Galvin, S.G., Miedzybrodzka, E.L., Leiter, A., Li, H.J., Glass, L.L., Ma, M.K.L., et al. (2019). Comparison of human and murine enteroendocrine cells by transcriptomic and peptidomic profiling. Diabetes 68, 1062–1072. 10.2337/db18-0883.

26. Haber, A.L., Biton, M., Rogel, N., Herbst, R.H., Shekhar, K., Smillie, C., Burgin, G., Delorey, T.M., Howitt, M.R., Katz, Y., et al. (2017). A single-cell survey of the small intestinal epithelium. Nature 551, 333–339. 10.1038/nature24489.

27. Billing, L.J., Larraufie, P., Lewis, J., Leiter, A., Li, J., Lam, B., Yeo, G.S., Goldspink, D.A., Kay, R.G., Gribble, F.M., and Reimann, F. (2019). Single cell transcriptomic profiling of large intestinal enteroendocrine cells in mice - Identification of selective stimuli for insulin-like peptide-5 and glucagon-like peptide-1 co-expressing cells. Mol Metab 29, 158–169. 10.1016/j.molmet.2019.09.001.

28. Gehart, H., van Es, J.H., Hamer, K., Beumer, J., Kretzschmar, K., Dekkers, J.F., Rios, A., and Clevers, H. (2019). Identification of enteroendocrine regulators by real-time single-cell differentiation mapping. Cell 176, 1158–1173.e1116. 10.1016/j.cell.2018.12.029.

29. Beumer, J., Puschhof, J., Bauza-Martinez, J., Martinez-Silgado, A., Elmentaite, R., James, K.R., Ross, A., Hendriks, D., Artegiani, B., Busslinger, G.A., et al. (2020). High-resolution mRNA and secretome atlas of human enteroendocrine cells. Cell 181, 1291–1306.e1219. 10.1016/j.cell.2020.04.036.

30. Basak, O., Beumer, J., Wiebrands, K., Seno, H., van Oudenaarden, A., and Clevers, H. (2017). Induced quiescence of Lgr5+ stem cells in intestinal organoids enables differentiation of hormone-producing enteroendocrine cells. Cell Stem Cell 20, 177–190 e174. 10.1016/j.stem.2016.11.001.

31. Wang, X., Yamamoto, Y., Wilson, L.H., Zhang, T., Howitt, B.E., Farrow, M.A., Kern, F., Ning, G., Hong, Y., Khor, C.C., et al. (2015). Cloning and variation of ground state intestinal stem cells. Nature 522, 173–178. 10.1038/nature14484.

32. Lopez-Diaz, L., Jain, R.N., Keeley, T.M., VanDussen, K.L., Brunkan, C.S., Gumucio, D.L., and Samuelson, L.C. (2007). Intestinal Neurogenin 3 directs differentiation of a bipotential secretory progenitor to endocrine cell rather than goblet cell fate. Dev Biol 309, 298–305. 10.1016/j.ydbio.2007.07.015.

33. Burclaff, J., Bliton, R.J., Breau, K.A., Ok, M.T., Gomez-Martinez, I., Ranek, J.S., Bhatt, A.P., Purvis, J.E., Woosley, J.T., and Magness, S.T. (2022). A proximal-to-distal survey of healthy adult human small intestine and colon epithelium by single-cell transcriptomics. Cell Mol Gastroenterol Hepatol 13, 1554–1589. 10.1016/j.jcmgh.2022.02.007.

34. McInnes, L., Healy, J., Saul, N., and Grossberger, L. (2018). UMAP: Uniform manifold approximation and projection. J Open Source Software 3, 861. 10.21105/joss.00861.

35. Fujita, Y., Chui, J.W., King, D.S., Zhang, T., Seufert, J., Pownall, S., Cheung, A.T., and Kieffer, T.J. (2008). Pax6 and Pdx1 are required for production of glucose-dependent insulinotropic polypeptide in proglucagon-expressing L cells. Am J Physiol Endocrinol Metab 295, E648–657. 10.1152/ajpendo.90440.2008.

36. Egerod, K.L., Engelstoft, M.S., Grunddal, K.V., Nohr, M.K., Secher, A., Sakata, I., Pedersen, J., Windelov, J.A., Fuchtbauer, E.M., Olsen, J., et al. (2012). A major lineage of enteroendocrine cells coexpress CCK, secretin, GIP, GLP-1, PYY, and neurotensin but not somatostatin. Endocrinology 153, 5782–5795. 10.1210/en.2012-1595.

37. Habib, A.M., Richards, P., Cairns, L.S., Rogers, G.J., Bannon, C.A., Parker, H.E., Morley, T.C., Yeo, G.S., Reimann, F., and Gribble, F.M. (2012). Overlap of endocrine hormone expression in the mouse intestine revealed by transcriptional profiling and flow cytometry. Endocrinology 153, 3054–3065. 10.1210/en.2011-2170.

38. Cao, J., Spielmann, M., Qiu, X., Huang, X., Ibrahim, D.M., Hill, A.J., Zhang, F., Mundlos, S., Christiansen, L., Steemers, F.J., et al. (2019). The single-cell transcriptional landscape of mammalian organogenesis. Nature 566, 496–502. 10.1038/s41586-019-0969-x.

39. Heintzman, N.D., Hon, G.C., Hawkins, R.D., Kheradpour, P., Stark, A., Harp, L.F., Ye, Z., Lee, L.K., Stuart, R.K., Ching, C.W., et al. (2009). Histone modifications at human enhancers reflect global cell-type-specific gene expression. Nature 459, 108–112. 10.1038/nature07829.

40. Buenrostro, J.D., Giresi, P.G., Zaba, L.C., Chang, H.Y., and Greenleaf, W.J. (2013). Transposition of native chromatin for fast and sensitive epigenomic profiling of open chromatin, DNA-binding proteins and nucleosome position. Nat Methods 10, 1213–1218. 10.1038/nmeth.2688.

41. Heumos, L., Schaar, A.C., Lance, C., Litinetskaya, A., Drost, F., Zappia, L., Lucken, M.D., Strobl, D.C., Henao, J., Curion, F., et al. (2023). Best practices for single-cell analysis across modalities. Nat Rev Genet 24, 550–572. 10.1038/s41576-023-00586-w.

42. Lynch, A.W., Theodoris, C.V., Long, H.W., Brown, M., Liu, X.S., and Meyer, C.A. (2022). MIRA: joint regulatory modeling of multimodal expression and chromatin accessibility in single cells. Nat Methods 19, 1097–1108. 10.1038/s41592-022-01595-z.

43. Solorzano-Vargas, R.S., Bjerknes, M., Wu, S.V., Wang, J., Stelzner, M., Dunn, J.C.Y., Dhawan, S., Cheng, H., Georgia, S., and Martin, M.G. (2019). The cellular regulators PTEN and BMI1 help mediate NEUROGENIN-3-induced cell cycle arrest. J Biol Chem 294, 15182–15192. 10.1074/jbc.RA119.008926.

44. Guillemot, F., Lo, L.C., Johnson, J.E., Auerbach, A., Anderson, D.J., and Joyner, A.L. (1993). Mammalian achaete-scute homolog 1 is required for the early development of olfactory and autonomic neurons. Cell 75, 463–476. 10.1016/0092-8674(93)90381-y.

45. Blaugrund, E., Pham, T.D., Tennyson, V.M., Lo, L., Sommer, L., Anderson, D.J., and Gershon, M.D. (1996). Distinct subpopulations of enteric neuronal progenitors defined by time of development, sympathoadrenal lineage markers and Mash-1-dependence. Development 122, 309–320. 10.1242/dev.122.1.309.

46. Pattyn, A., Simplicio,N., van Doorninck, J.H., Goridis, C., Guillemot, F., and Brunet, J.F. (2004). Ascl1/Mash1 is required for the development of central serotonergic neurons. Nat Neurosci 7, 589–595. 10.1038/nn1247.

47. Hatakeyama, J., Tomita, K., Inoue, T., and Kageyama, R. (2001). Roles of homeobox and bHLH genes in specification of a retinal cell type. Development 128, 1313–1322. 10.1242/dev.128.8.1313.

48. Imayoshi, I., Isomura, A., Harima, Y., Kawaguchi, K., Kori, H., Miyachi, H., Fujiwara, T., Ishidate, F., and Kageyama, R. (2013). Oscillatory control of factors determining multipotency and fate in mouse neural progenitors. Science 342, 1203–1208. 10.1126/science.1242366.

49. Sueda, R., and Kageyama, R. (2021). Oscillatory expression of Ascl1 in oligodendrogenesis. Gene Expr Patterns 41, 119198. 10.1016/j.gep.2021.119198.

50. Sueda, R., Imayoshi, I., Harima, Y., and Kageyama, R. (2019). High Hes1 expression and resultant Ascl1 suppression regulate quiescent vs. active neural stem cells in the adult mouse brain. Genes Dev 33, 511–523. 10.1101/gad.323196.118.

51. Bae, S., Bessho, Y., Hojo, M., and Kageyama, R. (2000). The bHLH gene Hes6, an inhibitor of Hes1, promotes neuronal differentiation. Development 127, 2933–2943. 10.1242/dev.127.13.2933.

52. Mastop, M., Bindels, D.S., Shaner, N.C., Postma, M., Gadella, T.W.J., Jr., and Goedhart, J. (2017). Characterization of a spectrally diverse set of fluorescent proteins as FRET acceptors for mTurquoise2. Sci Rep 7, 11999. 10.1038/s41598-017-12212-x.

53. Imayoshi, I., and Kageyama, R. (2014). bHLH factors in self-renewal, multipotency, and fate choice of neural progenitor cells. Neuron 82, 9–23. 10.1016/j.neuron.2014.03.018.

54. Borges, M., Linnoila, R.I., van de Velde, H.J., Chen, H., Nelkin, B.D., Mabry, M., Baylin, S.B., and Ball, D.W. (1997). An achaete-scute homologue essential for neuroendocrine differentiation in the lung. Nature 386, 852–855. 10.1038/386852a0.

55. Lanigan, T.M., DeRaad, S.K., and Russo, A.F. (1998). Requirement of the MASH-1 transcription factor for neuroendocrine differentiation of thyroid C cells. J Neurobiol 34, 126–134.

56. Ito, T., Udaka, N., Yazawa, T., Okudela, K., Hayashi, H., Sudo, T., Guillemot, F., Kageyama, R., and Kitamura, H. (2000). Basic helix-loop-helix transcription factors regulate the neuroendocrine differentiation of fetal mouse pulmonary epithelium. Development 127, 3913–3921. 10.1242/dev.127.18.3913.

57. Borromeo, M.D., Savage, T.K., Kollipara, R.K., He, M., Augustyn, A., Osborne, J.K., Girard, L., Minna, J.D., Gazdar, A.F., Cobb, M.H., and Johnson, J.E. (2016). ASCL1 and NEUROD1 reveal heterogeneity in pulmonary neuroendocrine tumors and regulate distinct genetic programs. Cell Rep 16, 1259–1272. 10.1016/j.celrep.2016.06.081.

58. Rudin, C.M., Poirier, J.T., Byers, L.A., Dive, C., Dowlati, A., George, J., Heymach, J.V., Johnson, J.E., Lehman, J.M., MacPherson, D., et al. (2019). Molecular subtypes of small cell lung cancer: a synthesis of human and mouse model data. Nat Rev Cancer 19, 289–297. 10.1038/s41568-019-0133-9.

59. Ireland, A.S., Micinski, A.M., Kastner, D.W., Guo, B., Wait, S.J., Spainhower, K.B., Conley, C.C., Chen, O.S., Guthrie, M.R., Soltero, D., et al. (2020). MYC drives temporal evolution of small cell lung cancer subtypes by reprogramming neuroendocrine fate. Cancer Cell 38, 60–78 e12. 10.1016/j.ccell.2020.05.001.

60. Nouruzi, S., Ganguli, D., Tabrizian, N., Kobelev, M., Sivak, O., Namekawa, T., Thaper, D., Baca, S.C., Freedman, M.L., Aguda, A., et al. (2022). ASCL1 activates neuronal stem cell-like lineage programming through remodeling of the chromatin landscape in prostate cancer. Nat Commun 13, 2282 10.1038/s41467-022-29963-5.

61. Cau, E., Gradwohl, G., Fode, C., and Guillemot, F. (1997). Mash1 activates a cascade of bHLH regulators in olfactory neuron progenitors. Development 124, 1611–1621. 10.1242/dev.124.8.1611.

62. Ma, Q., Chen, Z., del Barco Barrantes, I., de la Pompa, J.L., and Anderson, D.J. (1998). Neurogenin1 is essential for the determination of neuronal precursors for proximal cranial sensory ganglia. Neuron 20, 469–482. 10.1016/s0896-6273(00)80988-5.

63. Vierbuchen, T., Ostermeier, A., Pang, Z.P., Kokubu, Y., Sudhof, T.C., and Wernig, M. (2010). Direct conversion of fibroblasts to functional neurons by defined factors. Nature 463, 1035–1041. 10.1038/nature08797.

64. Chanda, S., Ang, C.E., Davila, J., Pak, C., Mall, M., Lee, Q.Y., Ahlenius, H., Jung, S.W., Sudhof, T.C., and Wernig, M. (2014). Generation of induced neuronal cells by the single reprogramming factor ASCL1. Stem Cell Reports 3, 282–296. 10.1016/j.stemcr.2014.05.020.

65. Wapinski, O.L., Vierbuchen, T., Qu, K., Lee, Q.Y., Chanda, S., Fuentes, D.R., Giresi, P.G., Ng, Y.H., Marro, S., Neff, N.F., et al. (2013). Hierarchical mechanisms for direct reprogramming of fibroblasts to neurons. Cell 155, 621–635. 10.1016/j.cell.2013.09.028.

66. Zhang, Y., Pak, C., Han, Y., Ahlenius, H., Zhang, Z., Chanda, S., Marro, S., Patzke, C., Acuna, C., Covy, J., et al. (2013). Rapid single-step induction of functional neurons from human pluripotent stem cells. Neuron 78, 785–798. 10.1016/j.neuron.2013.05.029.

67. Pang, Z.P., Yang, N., Vierbuchen, T., Ostermeier, A., Fuentes, D.R., Yang, T.Q., Citri, A., Sebastiano, V., Marro, S., Sudhof, T.C., and Wernig, M. (2011). Induction of human neuronal cells by defined transcription factors. Nature 476, 220–223. 10.1038/nature10202.

68. Raposo, A.A.S.F., Vasconcelos, F.F., Drechsel, D., Marie, C., Johnston, C., Dolle, D., Bithell, A., Gillotin, S., van den Berg, D.L.C., Ettwiller, L., et al. (2015). Ascl1 coordinately regulates gene expression and the chromatin landscape during neurogenesis. Cell Rep 10, 1544–1556. 10.1016/j.celrep.2015.02.025.

69. Soufi, A., Garcia, M.F., Jaroszewicz, A., Osman, N., Pellegrini, M., and Zaret, K.S. (2015). Pioneer transcription factors target partial DNA motifs on nucleosomes to initiate reprogramming. Cell 161, 555–568. 10.1016/j.cell.2015.03.017.

70. Pataskar, A., Jung, J., Smialowski, P., Noack, F., Calegari, F., Straub, T., and Tiwari, V.K. (2016). NeuroD1 reprograms chromatin and transcription factor landscapes to induce the neuronal program. EMBO J 35, 24–45. 10.15252/embj.201591206.

71. Matsuda, T., Irie, T., Katsurabayashi, S., Hayashi, Y., Nagai, T., Hamazaki, N., Adefuin, A.M.D., Miura, F., Ito, T., Kimura, H., et al. (2019). Pioneer Factor NeuroD1 Rearranges Transcriptional and Epigenetic Profiles to Execute Microglia-Neuron Conversion. Neuron 101, 472–485 e477. 10.1016/j.neuron.2018.12.010.

72. Paun, O., Tan, Y.X., Patel, H., Strohbuecker, S., Ghanate, A., Cobolli-Gigli, C., Llorian Sopena, M., Gerontogianni, L., Goldstone, R., Ang, S.L., et al. (2023). Pioneer factor ASCL1 cooperates with the mSWI/SNF complex at distal regulatory elements to regulate human neural differentiation. Genes Dev 37, 218–242. 10.1101/gad.350269.122.

73. Park, N.I., Guilhamon, P., Desai, K., McAdam, R.F., Langille, E., O’Connor, M., Lan, X., Whetstone, H., Coutinho, F.J., Vanner, R.J., et al. (2017). ASCL1 Reorganizes Chromatin to Direct Neuronal Fate and Suppress Tumorigenicity of Glioblastoma Stem Cells. Cell Stem Cell 21, 209–224 e207. 10.1016/j.stem.2017.06.004.

74. Kokubu, H., Ohtsuka, T., and Kageyama, R. (2008). Mash1 is required for neuroendocrine cell development in the glandular stomach. Genes Cells 13, 41–51. 10.1111/j.1365-2443.2007.01146.x.

75. Skene, P.J., Henikoff, J.G., and Henikoff, S. (2018). Targeted in situ genome-wide profiling with high efficiency for low cell numbers. Nat Protoc 13, 1006–1019. 10.1038/nprot.2018.015.

76. Zhu, Q., Liu, N., Orkin, S.H., and Yuan, G.C. (2019). CUT&RUNTools: a flexible pipeline for CUT&RUN processing and footprint analysis. Genome Biol 20, 192. 10.1186/s13059-019-1802-4.

77. Zaret, K.S. (2020). Pioneer transcription factors initiating gene network changes. Annu Rev Genet, 367–385. 10.1146/annurev-genet-030220-015007.

78. Sharma, A., Moore, M., Marcora, E., Lee, J.E., Qiu, Y., Samaras, S., and Stein, R. (1999). The NeuroD1/BETA2 sequences essential for insulin gene transcription colocalize with those necessary for neurogenesis and p300/CREB binding protein binding. Mol Cell Biol 19, 704–713. 10.1128/MCB.19.1.704.

79. Azzarelli, R., Hurley, C., Sznurkowska, M.K., Rulands, S., Hardwick, L., Gamper, I., Ali, F., McCracken, L., Hindley, C., McDuff, F., et al. (2017). Multi-site Neurogenin3 phosphorylation controls pancreatic endocrine differentiation. Dev Cell 41, 274–286 e275. 10.1016/j.devcel.2017.04.004.

80. Najle, S.R., Grau-Bove, X., Elek, A., Navarrete, C., Cianferoni, D., Chiva, C., Canas-Armenteros, D., Mallabiabarrena, A., Kamm, K., Sabido, E., et al. (2023). Stepwise emergence of the neuronal gene expression program in early animal evolution. Cell 186, 4676–4693 e4629. 10.1016/j.cell.2023.08.027.

81. Solorzano-Vargas, R.S., Bjerknes, M., Wang, J., Wu, S.V., Garcia-Careaga, M.G., Pitukcheewanont, P., Cheng, H., German, M.S., Georgia, S., and Martin, M.G. (2020). Null mutations of NEUROG3 are associated with delayed-onset diabetes mellitus. JCI Insight 5, e127657. 10.1172/jci.insight.127657.

82. Granja, J.M., Corces, M.R., Pierce, S.E., Bagdatli, S.T., Choudhry, H., Chang, H.Y., and Greenleaf, W.J. (2021). ArchR is a scalable software package for integrative single-cell chromatin accessibility analysis. Nat Genet 53, 403–411. 10.1038/s41588-021-00790-6.

83. Quinlan, A.R., and Hall, I.M. (2010). BEDTools: a flexible suite of utilities for comparing genomic features. Bioinformatics 26, 841–842. 10.1093/bioinformatics/btq033.

84. Langmead, B., and Salzberg, S.L. (2012). Fast gapped-read alignment with Bowtie 2. Nat Methods 9, 357–359. 10.1038/nmeth.1923.

85. Dibner, C., Sage, D., Unser, M., Bauer, C., d’Eysmond, T., Naef, F., and Schibler, U. (2009). Circadian gene expression is resilient to large fluctuations in overall transcription rates. EMBO J 28, 123–134. 10.1038/emboj.2008.262.

86. Sage, D., Unser, M., Salmon, P., and Dibner, C. (2010). A software solution for recording circadian oscillator features in time-lapse live cell microscopy. Cell Div 5, 17. 10.1186/1747-1028-5-17.

87. Ramirez, F., Ryan, D.P., Gruning, B., Bhardwaj, V., Kilpert, F., Richter, A.S., Heyne, S., Dundar, F., and Manke, T. (2016). deepTools2: a next generation web server for deep-sequencing data analysis. Nucleic Acids Res 44, W160–165. 10.1093/nar/gkw257.

88. Love, M.I., Huber, W., and Anders, S. (2014). Moderated estimation of fold change and dispersion for RNA-seq data with DESeq2. Genome Biol 15, 550. 10.1186/s13059-014-0550-8.

89. Wickham, H. (2016). ggplot2: Elegant Graphics for Data Analysis (Springer-Verlag New York).

90. Heinz, S., Benner, C., Spann, N., Bertolino, E., Lin, Y.C., Laslo, P., Cheng, J.X., Murre, C., Singh, H., and Glass, C.K. (2010). Simple combinations of lineage-determining transcription factors prime cis-regulatory elements required for macrophage and B cell identities. Mol Cell 38, 576–589. 10.1016/j.molcel.2010.05.004.

91. Pinello, L., Xu, J., Orkin, S.H., and Yuan, G.C. (2014). Analysis of chromatin-state plasticity identifies cell-type-specific regulators of H3K27me3 patterns. Proc Natl Acad Sci USA 111, E344–353. 10.1073/pnas.1322570111.

92. Schindelin, J., Arganda-Carreras, I., Frise, E., Kaynig, V., Longair, M., Pietzsch, T., Preibisch, S., Rueden, C., Saalfeld, S., Schmid, B., et al. (2012). Fiji: an open-source platform for biological-image analysis. Nat Methods 9, 676–682. 10.1038/nmeth.2019.

93. Robinson, J.T., Thorvaldsdottir, H., Winckler, W., Guttman, M., Lander, E.S., Getz, G., and Mesirov, J.P. (2011). Integrative genomics viewer. Nat Biotechnol 29, 24–26. 10.1038/nbt.1754.

94. Zhang, Y., Liu, T., Meyer, C.A., Eeckhoute, J., Johnson, D.S., Bernstein, B.E., Nusbaum, C., Myers, R.M., Brown, M., Li, W., and Liu, X.S. (2008). Model-based analysis of ChIP-Seq (MACS). Genome Biol 9, R137. 10.1186/gb-2008-9-9-r137.

95. R Core Team (2019). R: A Language and Environment for Statistical Computing (R Foundation for Statistical Computing).

96. Meers, M.P., Tenenbaum, D., and Henikoff, S. (2019). Peak calling by Sparse Enrichment Analysis for CUT&RUN chromatin profiling. Epigenetics Chromatin 12, 42. 10.1186/s13072-019-0287-4.

97. Hao, Y., Hao, S., Andersen-Nissen, E., Mauck, W.M., 3rd, Zheng, S., Butler, A., Lee, M.J., Wilk, A.J., Darby, C., Zager, M., et al. (2021). Integrated analysis of multimodal single-cell data. Cell 184, 3573–3587 e3529. 10.1016/j.cell.2021.04.048.

98. Stuart, T., Butler, A., Hoffman, P., Hafemeister, C., Papalexi, E., Mauck, W.M., 3rd, Hao, Y., Stoeckius, M., Smibert, P., and Satija, R. (2019). Comprehensive Integration of Single-Cell Data. Cell 177, 1888–1902 e1821. 10.1016/j.cell.2019.05.031.

99. Street, K., Risso, D., Fletcher, R.B., Das, D., Ngai, J., Yosef, N., Purdom, E., and Dudoit, S. (2018). Slingshot: cell lineage and pseudotime inference for single-cell transcriptomics. BMC Genomics 19, 477. 10.1186/s12864-018-4772-0.

100. Dobin, A., Davis, C.A., Schlesinger, F., Drenkow, J., Zaleski, C., Jha, S., Batut, P., Chaisson, M., and Gingeras, T.R. (2013). STAR: ultrafast universal RNA-seq aligner. Bioinformatics 29, 15–21. 10.1093/bioinformatics/bts635.

101. Cornwell, M., Vangala, M., Taing, L., Herbert, Z., Koster, J., Li, B., Sun, H., Li, T., Zhang, J., Qiu, X., et al. (2018). VIPER: Visualization Pipeline for RNA-seq, a Snakemake workflow for efficient and complete RNA-seq analysis. BMC Bioinformatics 19, 135. 10.1186/s12859-018-2139-9.

102. Huang, X., Gu, W., Zhang, J., Lan, Y., Colarusso, J.L., Li, S., Pertl, C., Lu, J., Kim, H., Zhu, J., et al. (2023). Stomach-derived human insulin-secreting organoids restore glucose homeostasis. Nat Cell Biol 25, 778–786. 10.1038/s41556-023-01130-y.

103. Corces, M.R., Trevino, A.E., Hamilton, E.G., Greenside, P.G., Sinnott-Armstrong, N.A., Vesuna, S., Satpathy, A.T., Rubin, A.J., Montine, K.S., Wu, B., et al. (2017). An improved ATAC-seq protocol reduces background and enables interrogation of frozen tissues. Nat Methods 14, 959–962. 10.1038/nmeth.4396.

104. Jadhav, U., Cavazza, A., Banerjee, K.K., Xie, H., O’Neill, N.K., Saenz-Vash, V., Herbert, Z., Madha, S., Orkin, S.H., Zhai, H., and Shivdasani, R.A. (2019). Extensive recovery of embryonic enhancer and gene memory stored in hypomethylated enhancer DNA. Mol Cell 74, 542–554 e545. 10.1016/j.molcel.2019.02.024.

105. Hainer, S.J., and Fazzio, T.G. (2019). High-Resolution Chromatin Profiling Using CUT&RUN. Curr Protoc Mol Biol 126, e85. 10.1002/cpmb.85.

106. Feng, J., Liu, T., Qin, B., Zhang, Y., and Liu, X.S. (2012). Identifying ChIP-seq enrichment using MACS. Nat Protoc 7, 1728–1740. 10.1038/nprot.2012.101.

107. Stuart, T., Srivastava, A., Madad, S., Lareau, C.A., and Satija, R. (2021). Single-cell chromatin state analysis with Signac. Nat Methods 18, 1333–1341. 10.1038/s41592-021-01282-5.

108. Clement, K., Rees, H., Canver, M.C., Gehrke, J.M., Farouni, R., Hsu, J.Y., Cole, M.A., Liu, D.R., Joung, J.K., Bauer, D.E., and Pinello, L. (2019). CRISPResso2 provides accurate and rapid genome editing sequence analysis. Nat Biotechnol 37, 224–226. 10.1038/s41587-019-0032-3.

109. Pliner, H.A., Packer, J.S., McFaline-Figueroa, J.L., Cusanovich, D.A., Daza, R.M., Aghamirzaie, D., Srivatsan, S., Qiu, X., Jackson, D., Minkina, A., et al. (2018). Cicero predicts cis-regulatory DNA interactions from single-cell chromatin accessibility data. Mol Cell 71, 858–871 e858. 10.1016/j.molcel.2018.06.044.

